# Multi-omics reveals accumulation mechanisms underlying differences in sesquiterpenoid content across different growth environments of *Atractylodes macrocephala* rhizome

**DOI:** 10.64898/2026.10.10.758175

**Authors:** Meili Chen, Shiqi Yu, Yakun Wang, Yi Zheng, Rongxin Zou, Juan Liang, Wentao Zhu, Wei Fang, Dianlei Wang, Nianxia Sun

## Abstract

This study aimed to explore mechanisms driving differential sesquiterpenoid accumulation in *Atractylodes macrocephala* rhizome (AMR) under different growth environments. Root and rhizosphere soil samples were collected from wild, imitated wild, one- and two-year cultivated AMR. Combined metabolomic, transcriptomic and microbiomic profiling was carried out to investigate alterations in sesquiterpenoid levels, root-associated microbiota, and terpenoid biosynthetic genes under different growth environments. Metabolomics revealed that wild AMR exhibited higher pyruvate consumption and accumulated more atractylone and atractylenolides. Transcriptomic analysis demonstrated that the genes encoding key MEP pathway enzymes (DXS, DXR, MCT, HDS) and the genes encoding the sesquiterpenoid biosynthetic enzyme GDS were significantly upregulated in wild AMR. In contrast, the genes encoding triterpenoid synthetic enzymes (SQS, SQE, β-AS) were downregulated, redirecting metabolic precursors toward sesquiterpenoid synthesis. Microbial community structures differed substantially across groups. Further analysis suggested rhizosphere microorganisms were important contributors to AMR medicinal quality, and sesquiterpenoid metabolites accumulated in AMR roots, together with the rhizosphere microenvironment, jointly shape specific rhizosphere fungal assembly. This study systematically revealed the synergistic regulatory relationships among sesquiterpenoids metabolism, gene expression, growth environment, and root-associated microbes, providing multi-omics evidence for clarifying the mechanism by which these factors coordinately regulate the quality of AMR.

## Introduction

*Atractylodes macrocephala* rhizome (AMR), a perennial species of the Compositae family, is a traditional Chinese medicinal plant. It is first documented in *Shennong’s Classic of Materia Medica* and classified as a top-grade medicinal herb. With therapeutic actions of spleen invigoration, qi replenishment, dampness elimination, and diuresis promotion, it is revered as the foremost medicinal herb for spleen tonification. Its main active components are sesquiterpenoids, which exhibit anti-inflammatory and antioxidant activities (Qu et al., 2024). The primary production regions of AMR are concentrated in Zhejiang and Anhui Provinces. Specifically, wild AMR, a variety endemic to the Qimen Mountain area, is a renowned genuine medicinal material of Anhui Province. Unfortunately, it is difficult to acquire wild AMR since it has become endangered due to environmental challenges and overexploitation. With the application of artificial cultivation technology, AMR cultivated in Bozhou City, Anhui Province, is one of the most artificially grown herbs of AMR. Artificially cultivated AMR exhibits several issues, including reduced drug efficacy, impaired biosynthetic capacity of sesquiterpenoid bioactive components, and significantly reduced contents of effective ingredients (Zhou et al., 2025). To improve the quality of cultivated AMR, the imitated wild cultivation approach is adopted. Nevertheless, sesquiterpenoid contents exhibit substantial variation driven by geographical origin and growth conditions, exerting profound effects on the medicinal quality of the herb. Mechanisms responsible for differential sesquiterpenoid accumulation are still incompletely characterised.

As plant secondary metabolites (PSMs), sesquiterpenoids are dispensable for plant growth and development, yet they serve as the fundamental chemical basis for the clinical efficacy of medicinal plants. Their synthesis and accumulation are usually closely associated with plant adaptation to abiotic stresses (such as drought and pest infestation), participation in signal transduction pathways, and ecological interactions with the environment (Pant et al., 2021). Sesquiterpenoids are derived from the five-carbon building blocks isopentenyl diphosphate (IPP) and dimethylallyl diphosphate (DMAPP), which are biosynthesized via the mevalonic acid (MVA) and 2-C-methyl-D-erythritol 4-phosphate (MEP) pathways. Transcription factors can regulate the biosynthesis of these specialized metabolites by tightly controlling the expression of biosynthetic genes, forming complex transcriptional regulatory networks. Transcriptome analysis represents a core approach for screening regulatory genes and elucidating underlying metabolic pathways (Shi et al., 2024). For example, Zhao et al. (2021) performed comparative transcriptome analysis on the rhizomes of one-year-old, two-year-old, and three-year-old *Atractylodes chinensis*, successfully screening 52 differentially expressed genes related to sesquiterpenoid and triterpenoid biosynthesis and validated the expression patterns of key enzyme genes (Zhao et al., 2021). Based on this, the present study intended to employ transcriptome analysis to screen differentially expressed genes related to the biosynthesis and accumulation of sesquiterpenoids in different AMR samples, thereby providing insights into how environmental factors modulate secondary metabolite accumulation in AMR.

With advances in high-throughput sequencing and microbiome technologies, microorganisms have been increasingly recognized for their ability to regulate PSMs. The important roles and regulatory mechanisms of endophytes, rhizosphere microorganisms, and their communities in the accumulation of PSMs are mainly achieved by regulating the expression of key genes in PSMs, secreting phytohormones or effector molecules, and mediating signal pathway crosstalk. They form a bidirectional interaction network with PSMs, jointly influencing plant metabolic phenotypes and environmental adaptability (Pang et al., 2021). In the context of sesquiterpenoids biosynthesis and accumulation, microorganisms dominated by endophytic bacteria and plant growth-promoting rhizobacteria (mainly *Bacillus*, *Pseudomonas*, and *Acinetobacter*) have been shown to promote plant growth and the synthesis of sesquiterpenoid metabolites (Vaghela & Gohel, 2022; Wang et al., 2014; Yang et al., 2019). Additionally, fungi, mainly root endophytic fungi, arbuscular mycorrhizal fungi, and even pathogenic fungi, also played important roles in promoting plant growth and sesquiterpenoid accumulation (Lv et al., 2024). Different microbial groups could function individually or jointly, providing the basis for the microbial-assisted production of sesquiterpenoids. Nevertheless, the association between AMR and microorganisms was highly complex and the underlying mechanisms had not yet been fully elucidated.

This study adopted a multi-omics approach integrating metabolomics, transcriptomics and microbiome analyses to establish a comprehensive analytical framework. First, combined transcriptomic and metabolomic analyses were performed to systematically dissect the molecular mechanism underlying differential accumulation of sesquiterpenoid active constituents in AMR under different growth environments, and to identify the key metabolic pathways and core enzyme genes governing sesquiterpenoid biosynthesis. On this basis, by incorporating soil physicochemical properties and rhizosphere microbiome data, we further revealed the regulatory effects of soil conditions and microbial communities on sesquiterpenoid accumulation in AMR, anddemonstratedthatrhizospheremicroorganismsservedasthe keyfactormediatingqualityvariations ofAMR across diverse habitats. Finally, multi-omics correlation analysis further clarified the influences of key microbial taxa on sesquiterpenoid contents and the expression of key biosynthetic genes in the sesquiterpenoid pathway of AMR. This study aimed to establish a theoretical foundation for improving the medicinal quality of AMR medicinal materials.

## Materials and methods

### 2.1. Material collection

Wild AMR specimens were collected from the primary growing regions in the Qimen Mountains, Huangshan City, Anhui Province (Wild: 29.9711° N, 117.2200° E). Imitated wild AMR specimens were collected from the Dabie Mountains, Lu’an City, Anhui Province (ImW: 29.9851° N, 117.2035° E). One-year-old artificially cultivated AMR and two-year-old artificially cultivated AMR specimens were collected from Qiaocheng District, Bozhou City (C1Y and C2Y: 33.8756° N, 115.7823° E). Samples were authenticated by Professor Peng Huang (Anhui University of Chinese Medicine). Wild and imitated wild AMR were excavated from field soil, and loosely attached particles were removed by gentle shaking. Rhizosphere soil within 1 mm of root surfaces was isolated by phosphate buffer washing. Two sonication steps were applied to roots to strip surface-associated microorganisms, followed by sample storage at −80 °C.

### 2.2. Metabolomic profiling and analysis

Samples were weighed, snap-frozen in liquid nitrogen and stored at −80 °C. After grinding in liquid nitrogen, 500 mg sample was added into a 20 mL headspace vial containing 2 mL saturated NaCl solution to suppress enzymatic activity and sealed with TFE-silicone septum crimp caps. Following published SPME protocols (Yuan et al., 2022; Yuan et al., 2024), samples were incubated at 60 °C for 5 min, and headspace extraction was conducted with a 120 µm DVB/CWR/PDMS SPME Arrow (60 °C, 15 min). Volatiles were desorbed at 250 °C for 5 min in the GC inlet. For untargeted metabolomics, metabolite profiling was determined by gas chromatography-tandem time-of-flight mass spectrometry (GC-TOFMS) with full-scan acquisition. Global volatile fingerprints and tentative metabolite annotation were performed based on the NIST spectral library. For widely targeted metabolomics, samples were analyzed on an Agilent 8890 GC-7000D MS system in SIM mode. Characteristic ions and retention parameters were predefined based on an in-house volatile metabolite library for accurate quantification of known volatile compounds. The GC column was DB-5MS (30 m × 0.25 mm × 0.25 μm) with helium carrier gas at 1.2 mL/min and an injector temperature of 250 °C. The oven temperature was held at 40 °C for 3.5 min, ramped to 100 °C, 180 °C and 280 °C at 10 °C/min, 7 °C/min and 25 °C/min, respectively, and maintained at 280 °C for 5 min. Under 70 eV EI ionization, the quadrupole, ion source and transfer line temperatures were set at 150 °C, 230 °C and 280 °C, respectively. Unsupervised PCA was implemented in R after unit-variance scaling. Differential metabolites were defined as VIP > 1 and |log₂FC| ≥ 1.0. OPLS-DA models were constructed with MetaboAnalystR (Thévenot et al., 2015). Data were log₂-transformed and mean-centered before modelling, and 200 permutations were used to exclude overfitting. Metabolites were annotated against the KEGG Compound database and assigned to KEGG Pathways.

### 2.3 Determination of sesquiterpenoid contents by HPLC

After excess soil was removed, AMR roots were washed, dried, sliced, ground, and passed through an 80-mesh sieve. Sample (2 g) was extracted with 10 mL methanol by ultrasonication (60 Hz, 250 W, 60 min), centrifuged (3500 rpm, 10 min), and filtered (0.22 μm). Atractylenolide I, II, III, and atractylon were quantified by HPLC (Thermo Scientific UltiMate 3000; Dikma Kromasil C18, 250 × 4.6 mm, 5 μm) with acetonitrile–water gradient elution at 1.0 mL/min, 30 °C, and detection at 220 and 276 nm (10 μL injection). (Wang et al., 2026).

### 2.4. Transcriptome sequencing and data analysis

TRIzol Reagent (Invitrogen Life Technologies) was used to isolate total RNA from four AMR sample sets. The quality metrics including RNA concentration, purity and integrity were determined on the Agilent 2100 Bioanalyzer. Three micrograms of RNA were used for RNA sample preparation and sequencing library construction. Library quantification was performed using a Qubit system. Following library construction, paired-end 150 (PE150) sequencing was carried out on a DNBSEQ-T7 platform at Bioyi Biotechnology Co., Ltd. (Wuhan, China).

Raw reads were processed to yield clean reads, and Trinity was applied for de novo transcriptome assembly. Transcript abundance was quantified and normalized to fragments per kilobase of transcript per million mapped reads (FPKM). Differentially expressed genes (DEGs) were identified with the screening criteria of |log₂Fold Change| > 1.2 and *P* < 0.05. KEGG pathway enrichment analysis was conducted using the clusterProfiler software, with *P* < 0.05 set as the threshold for significant enrichment.

### 2.5. Soil physicochemical analyses

Soil physicochemical properties were analyzed using standard methods. Soil pH was measured in a 1:2.5 (w/v) soil-deionized water suspension after 1.5 h of stabilization using a calibrated OHA-2F pH meter. Soil organic matter was determined by potassium dichromate oxidation, with organic carbon quantified via ferrous sulfate titration after high-temperature digestion. Total nitrogen was measured by the Kjeldahl method through concentrated sulfuric acid digestion, alkali distillation, boric acid absorption, and hydrochloric acid titration, with blank tests included. Alkali-hydrolyzable nitrogen was determined by alkaline diffusion after 24 h incubation at 40 °C with zinc-ferrous sulfate reduction. Available phosphorus was extracted with sodium bicarbonate solution under constant shaking, followed by color reaction and absorbance detection at 880 nm. All measurements were performed in three biological replicates (Zuo et al., 2026).

### 2.6. Sequencing and annotation of endophytic and rhizosphere bacteria and fungi

Total genomic DNA was extracted from root and rhizosphere soil samples using the VAMNE Stool/Soil DNA Extraction Kit (Vazyme #DM401-C3). Agarose gel electrophoresis and NanoDrop NC2000 spectrophotometry (Thermo Fisher Scientific, USA) were applied to examine DNA purity and concentration. Targeted PCR amplification was separately performed for bacterial 16S rRNA and fungal ITS regions using sample-specific primer pairs appended with 7-bp barcodes to support multiplex sequencing. The bacterial V3–V4 region was amplified with the forward primer 338F (5’-ACTCCTACGGGAGGCAGCA-3’) and reverse primer 806R (5’-GGACTACHVGGGTWTCTAAT-3’), whereas the fungal ITS region was amplified using ITS1F (5’-CTTGGTCATTTAGAGGAAGTAA-3’) and ITS2R (5’-GCTGCGTTCTTCATCGATGC-3’).

All PCRs were performed in a 25 μL uniform reaction system: 5 μL 5× reaction buffer, 0.25 μL Fast pfu DNA Polymerase, 2 μL 2.5 mM dNTP mixture, 1 μL of each 10 μM primer, 1 μL template DNA, and 14.75 μL double-distilled water. The thermal program comprised an initial denaturation at 98 °C for 5 min, 25 cycles (98 °C for 30 s, 53 °C for 30 s, 72 °C for 45 s), and a final extension at 72 °C for 5 min. PCR amplicons were purified with Vazyme VAHTSTM DNA Clean Beads and quantified using the Quant-iT PicoGreen dsDNA Assay Kit (Invitrogen, USA). Purified qualified amplicons were pooled equimolarly and subjected to paired-end 2 × 250 bp sequencing on the Illumina MiSeq platform (MiSeq Reagent Kit v3) at Bioyi Biotechnology Co., Ltd. (Wuhan, China).

### 2.7. Microbial data analysis

α-diversity was assessed using the Shannon, Simpson, Chao1, and Observed species indices. Bacterial 16S sequences were taxonomically assigned into ASVs using the classify-sklearn naïve Bayes classifier (feature-classifier plugin) against the Greengenes 2 Database, whereas taxonomic annotation of fungal ITS sequences was performed based on the UNITE database. β-diversity was estimated by PCoA based on the Bray–Curtis metric. Genus-level bacterial and fungal biomarkers with significant variation across groups were identified using the LEfSe algorithm (|LDA score| > 2).

### 2.8. Statistical analysis

Metabolomic, transcriptomic, and microbiomic sequencing were all performed with three biological replicates. Heatmap analysis of transcriptomic gene expression was performed using the bio-cloud, a comprehensive platform for biological data analysis and visualization (https://bio-cloud.aptbiotech.com/plus/#/tools). Values were presented as mean ± standard deviation (SD) (n ≥ 3). Variations in atractylon and atractylenolide I, II, and III concentrations across four AMR groups were evaluated by one-way ANOVA, with the significance threshold set at *P* < 0.05. All statistical computations were performed in IBM SPSS Statistics 26.0. Weighted gene co-expression network analysis (WGCNA) was applied to screen core genes highly correlated with key sesquiterpenoid metabolites. Spearman correlation coefficients among microbial OTUs/ASVs, transcript levels and metabolite contents were calculated using the Metabo-Cloud platform for the construction of correlation networks. Correlation statistics and visualization were implemented in Cytoscape (v3.10). The *P* values of Spearman correlation analyses were adjusted by the Benjamini–Hochberg method, and FDR < 0.05 was regarded as the threshold for statistical significance.

## Result

### 3.1 Qualitative and quantitative analysis of the metabolites

To investigate the influence of different producing areas and growth years on the chemical components of AMR, in this study, four types of AMR samples were selected, namely wild AMR, imitated wild AMR, one-year-old artificially cultivated AMR, and two-year-old artificially cultivated AMR, and untargeted metabolomics were determined by gas-chromatography-tandem time-of-flight mass spectrometry (GC-TOFMS), and widely-targeted metabolomics was detected using gas chromatography-mass spectrometry (GC-MS). A total of 577 metabolites were detected via untargeted metabolomics analysis (Table S1). Among these, organic acids and derivatives represented the predominant class in all samples, comprising 28.23%. (Fig. S1A). A total of 40, 70, 28, and 39 differential metabolites were screened from the four comparison groups (ImW vs Wild, C2Y vs Wild, C2Y vs ImW, and C1Y vs C2Y), respectively (Fig. S1B). Metabolic pathway enrichment analysis was implemented to uncover pathways related to metabolites with significant abundance variations. (Fig. S1C-F). The results revealed that multiple metabolic pathways were significantly enriched across the four comparison groups, including glycolysis/gluconeogenesis, pyruvate metabolism, the citrate cycle (TCA cycle), the pentose phosphate pathway, and carbon metabolism. (*P* < 0.05). D-fructose-6-P exhibited significantly increased abundances in three comparison groups: C2Y vs ImW, C2Y vs Wild, and C2Y vs C1Y, whereas pyruvate was remarkably decreased in ImW vs Wild ( *P* < 0.05) (Table S2). As a common intermediate for glycolysis and the pentose-phosphate pathway, D-fructose-6-P was derived from β-D-glucose-6-phosphate catalyzed by glucose-6-phosphate isomerase in the pentose-phosphate pathway. Subsequently, pyruvate was produced via glycolysis, which served as the key carbon precursor for plant sesquiterpenoid biosynthesis. The joint effects of the two pathways provided precursors and energy for the biosynthesis of sesquiterpenoids.

For widely-targeted metabolomics data, metabolite profiling of the AMR samples resulted in the identification of 746 metabolites. (Table S3). Among these, Terpenoids had the highest proportion in all samples, accounting for 31.01% (Fig. 1A). The results of Principal Component Analysis (PCA) indicated that the metabolite profiles of different varieties were effectively separated (Fig. S2). The distinct separation of samples from different samples revealed significant variability in the metabolites. For comparison of the metabolic differences among groups, Variable Importance in Projection (VIP) values derived from the OPLS-DA model, combined with fold change (FC) values, were used to screen differential metabolites from the annotated metabolite data. Metabolites that met the threshold of VIP > 1 together with either FC ≥ 2 or FC ≤ 0.5 were identified as significantly differential metabolites (Thévenot et al., 2015). A total of 505, 314, 262, and 329 differential metabolites were screened from the four comparison groups (ImW vs Wild, C2Y vs Wild, C2Y vs ImW, and C1Y vs C2Y), respectively (Fig. 1B). The differentially abundant metabolites were subjected to pathway enrichment analysis to identify their associated metabolic pathways. (Fig. 1C-F). The findings revealed that the sesquiterpenoid and triterpenoid biosynthetic pathway was significantly enriched in C2Y vs ImW group and C1Y vs C2Y group (*P* < 0.05).

**Fig. 1.**
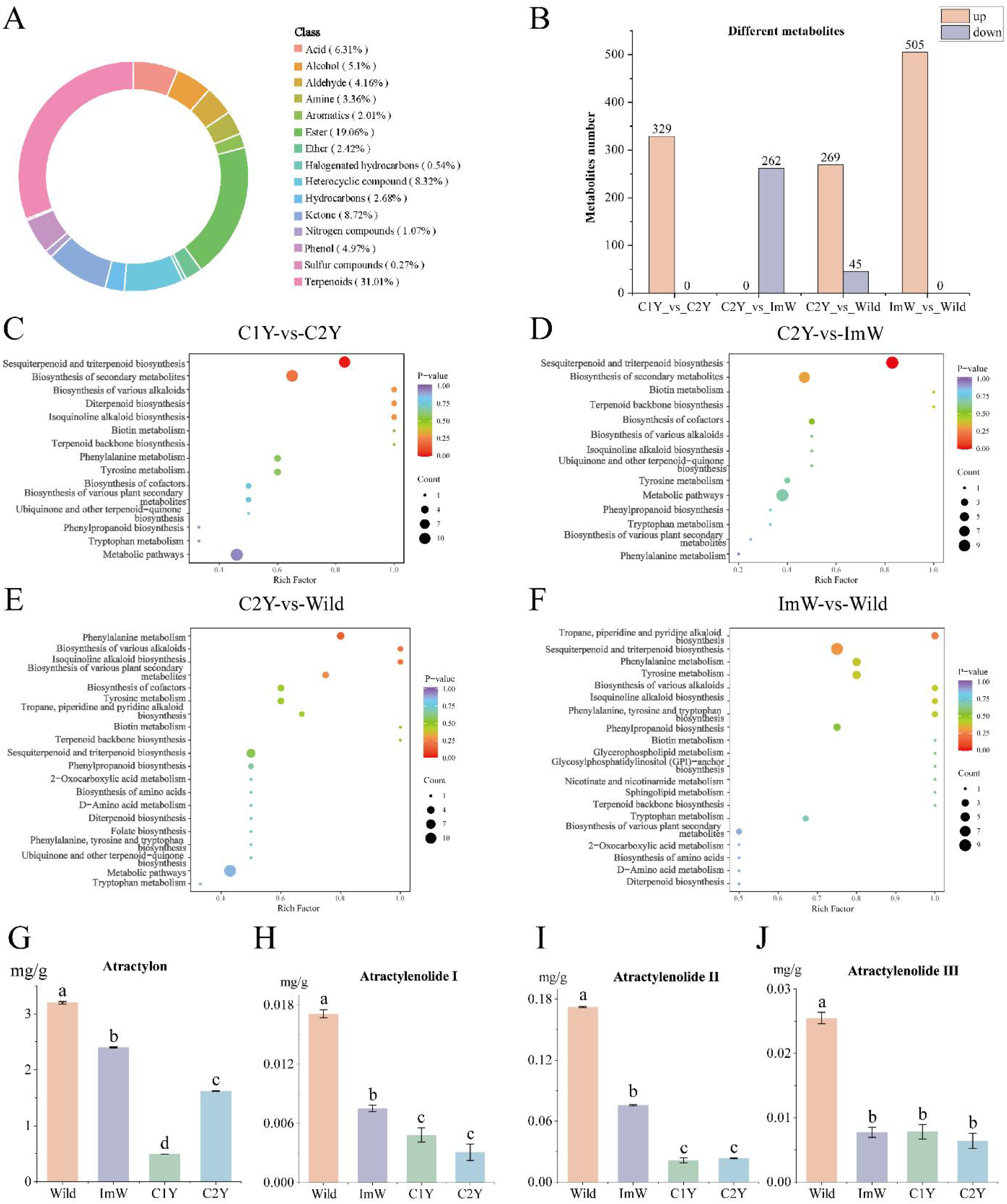
Metabolomic profiling and quantitative analysis of key bioactive components in four AMR sample groups. (A) The classification of all identified metabolites; (B) Differences in metabolite contents; (C-F) Pathway enrichment of differential metabolites. Bubble size and colour gradient represented the number of enriched differential metabolites and the enrichment *P*-value, respectively. (G–J) The contents of atractylon (G), atractylenolide Ⅰ (H), atractylenolide Ⅱ (I), and atractylenolide Ⅲ (J) in AMR. Data were shown as mean ± SD (n = 3), with distinct lowercase letters denoting significant differences at *P* < 0.05.

Sesquiterpenoids were the most important secondary metabolites in AMR. A total of 12 differential metabolites belonging to the sesquiterpenoid class were significantly enriched across the four comparison groups, as detailed in Table 1. Among them, (E, E)-Farnesol, (E, E)-α-Farnesol, (E, E)-β-Farnesol, (+)-α-Santalene, Valencene, Longifolene, α-Humulene, δ-Cadinene and β-Selinene were significantly upregulated in the C1Y vs. C2Y group. (E, E)-Farnesol, (E, E)-α-Farnesol, (E, E)-β-Farnesol, (+)-α-Santalene, Valencene, Longifolene, α-Humulene, δ-Cadinene and β-Selinene were significantly downregulated in the C2Y vs. ImW group. (E, E)-Farnesol, (S, E)-Nerolidol, (+)-α-Santalene and α-Humulene were significantly upregulated in the C2Y vs Wild group. Longifolene and β-Selinene were significantly downregulated in the C2Y vs Wild group. (E, E)-Farnesol, (E, E)-α-Farnesol, (E, E)-β-Farnesol, (+)-α-Santalene, Germacrene B, Valencene, Longifolene, α-Humulene and δ-Cadinene were significantly upregulated in the ImW vs. Wild group.

**Table 1.** Differential sesquiterpenoid metabolites informations of different types of AMR.

| Name | Exact mass | Chemical formula | PubChem | KEGG compound ID | CAS number |
| --- | --- | --- | --- | --- | --- |
| (E, E)-Farnesol | 222.1984 | C<sub>15</sub>H<sub>26</sub>O | 4357 | C01126 | 106-28-5 |
| (E, E)-α-Farnesol | 204.1878 | C<sub>15</sub>H<sub>24</sub> | 11855 | C09665 | 502-61-4 |
| (E, E)-β-Farnesol | 204.1878 | C<sub>15</sub>H<sub>24</sub> | 11856 | C09666 | 18794-84-8 |
| (+)-α-Santalene | 204.1878 | C<sub>15</sub>H<sub>24</sub> | 135626203 | C19736 | 512-61-8 |
| Germacrene B | 204.1878 | C<sub>15</sub>H<sub>24</sub> | 11862 | C09672 | 15423-57-1 |
| Valencene | 204.1878 | C<sub>15</sub>H<sub>24</sub> | 96023682 | C17277 | 4630-07-3 |
| Longifolene | 204.1878 | C<sub>15</sub>H<sub>24</sub> | 11889 | C09699 | 475-20-7 |
| α-Humulene | 204.1878 | C<sub>15</sub>H<sub>24</sub> | 11874 | C09684 | 6753-98-6 |
| δ-Cadinene | 204.1878 | C<sub>15</sub>H<sub>24</sub> | 8630 | C06394 | 483-76-1 |
| (S, E)-Nerolidol | 222.1984 | C<sub>15</sub>H<sub>26</sub>O | 11893 | C09704 | 7212-44-4 |
| β-Selinene | 204.1878 | C<sub>15</sub>H<sub>24</sub> | 11911 | C09723 | 17066-67-0 |
| (S)-β-Bisabolene | 204.1878 | C<sub>15</sub>H<sub>24</sub> | 96023299 | C16775 | 495-61-4 |

To further determine how several important sesquiterpenoids of AMR differ among the different groups, absolute quantification analysis of atractylone and atractylenolide (Ⅰ, II, III) were performed and the results were shown in Fig. 1G-J (Table S4). These four bioactive components were significantly more abundant in wild AMR than in imitated wild and artificially cultivated AMR. The imitated wild group had significantly higher contents of atractylone and atractylenolide (I, II) relative to artificially cultivated AMR. Two-year-old artificially cultivated AMR exhibited a significantly higher atractylone content than one-year-old artificially cultivated AMR (*P* < 0.05).

### 3.2 Transcriptome analysis of different varieties of AMR

To clarify whether distinct expression patterns of differential metabolite-associated genes existed across AMR varieties, the transcriptome of AMR was sequenced, assembled, and annotated (Table S5). PCA clearly separated the four AMR groups (Fig. 2A), indicating distinct gene expression differences among them. Genes with FC ≥ 1.2 and *P* ≤ 0.05 were considered significantly differentially expressed. In the pairwise comparisons among all groups, the comparison between wild and imitated wild AMR yielded the greatest number of differentially expressed genes, comprising 1,696 upregulated and 5,014 downregulated genes. By contrast, the comparison between one-year-old and two-year-old cultivated AMR produced the fewest DEGs, with 600 upregulated and 752 downregulated genes. The comparison group of two-year-old artificially cultivated AMR and wild AMR contained 922 upregulated genes and 3303 downregulated genes. The pairwise comparison between two-year-old artificially cultivated AMR and imitated wild AMR identified 756 upregulated and 752 downregulated genes (Fig. 2B). Kyoto Encyclopedia of Genes and Genomes (KEGG) pathway enrichment analysis revealed that these differentially expressed genes were mainly enriched in the following pathways: plant hormone signal transduction, MAPK signaling pathway – plant, flavonoid biosynthesis, photosynthesis – antenna proteins, stilbenoid, diarylheptanoid and gingerol biosynthesis (ImW vs Wild), phenylpropanoid biosynthesis, pentose and glucuronate interconversions (C2Y vs ImW), and photosynthesis (C1Y vs C2Y) (Fig. 2C-F). The differences in metabolite contents in AMR under different growth conditions may result from the enriched metabolic pathways of these differentially expressed genes.

**Fig. 2.**
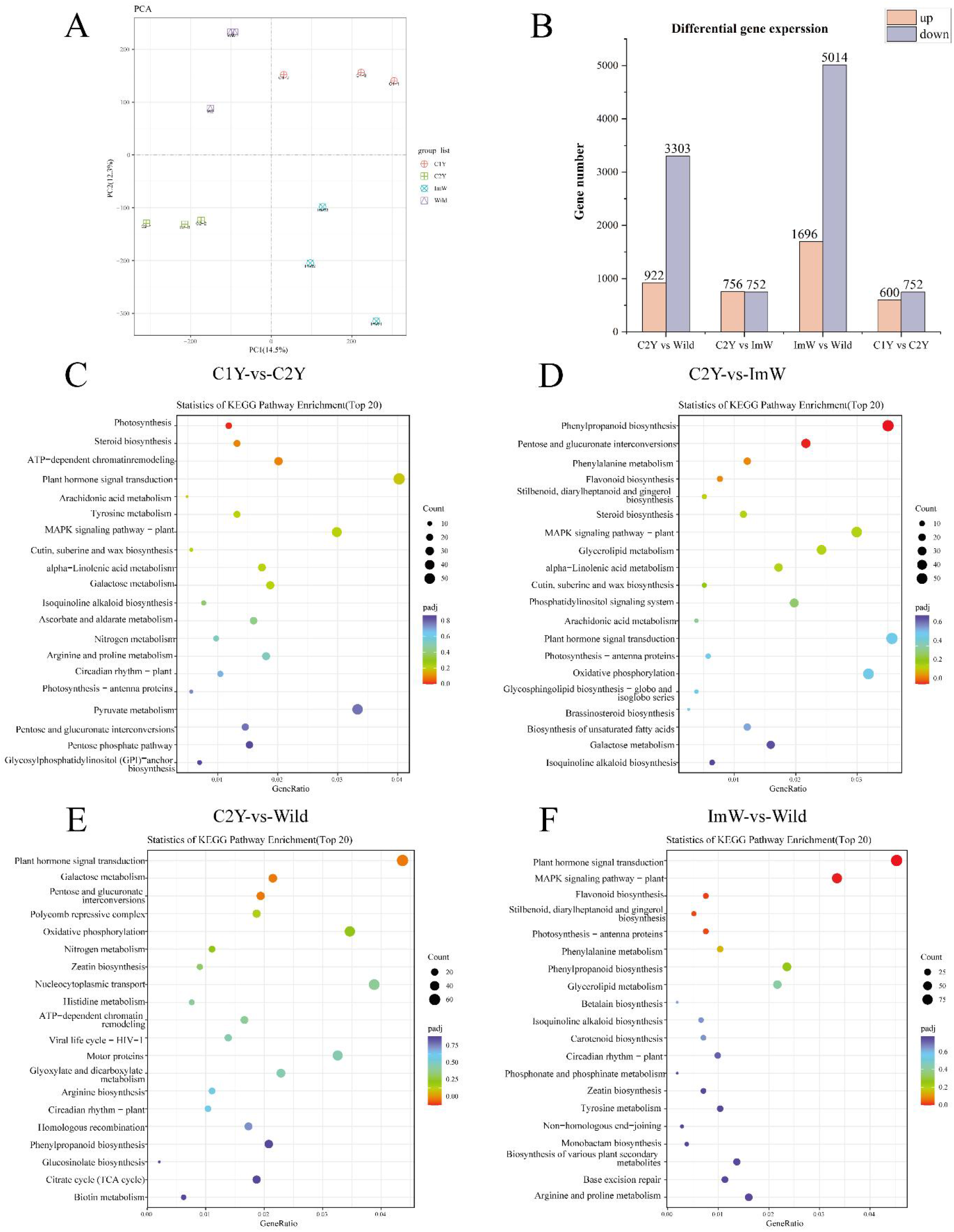
Gene expression patterns and DEG enrichment. (A) PCR results of gene expression; (B) Numbers of up-regulated and down-regulated differentially expressed genes in each comparison group; (C–F) Pathway enrichment of DEGs in each comparison group. Bubble size represents the number of enriched genes (Count), and the colour gradient indicates the enrichment *P*-value; Gene Ratio is the percentage of total DEGs in a given pathway term.

Plant hormone signal transduction pathways played a crucial role in regulating the growth, development, and secondary metabolism of medicinal plants (Alum et al., 2025). The jasmonic acid (JA) signaling pathway showed enrichment of DEGs among plant hormone signal transduction pathways. It has been shown that JA might serve as the core pathway governing the biosynthesis of sesquiterpenoid active ingredients (Chen et al., 2019). As shown in Fig. 3A, the gene encoding jasmonate-amino acid synthetase (JAR1) was upregulated in one-year-old artificially cultivated AMR, promoting the synthesis of active jasmonic acid (JA-Ile). The gene encoding coronatine-insensitive 1 (COI1) was significantly upregulated in wild AMR, while the genes encoding jasmonate ZIM-domain protein (JAZ) and myelocytomatosis transcription factor 2 (MYC2) were upregulated in both wild and simulated-wild samples. It was revealed that MYC2 could activate expression of the downstream target gene octadecanoid-responsive *catharanthus* AP2/ERF-domain protein 3 (ORCA3), which in turn regulated sesquiterpenoid biosynthesis (Montiel et al., 2011).

**Fig. 3.**
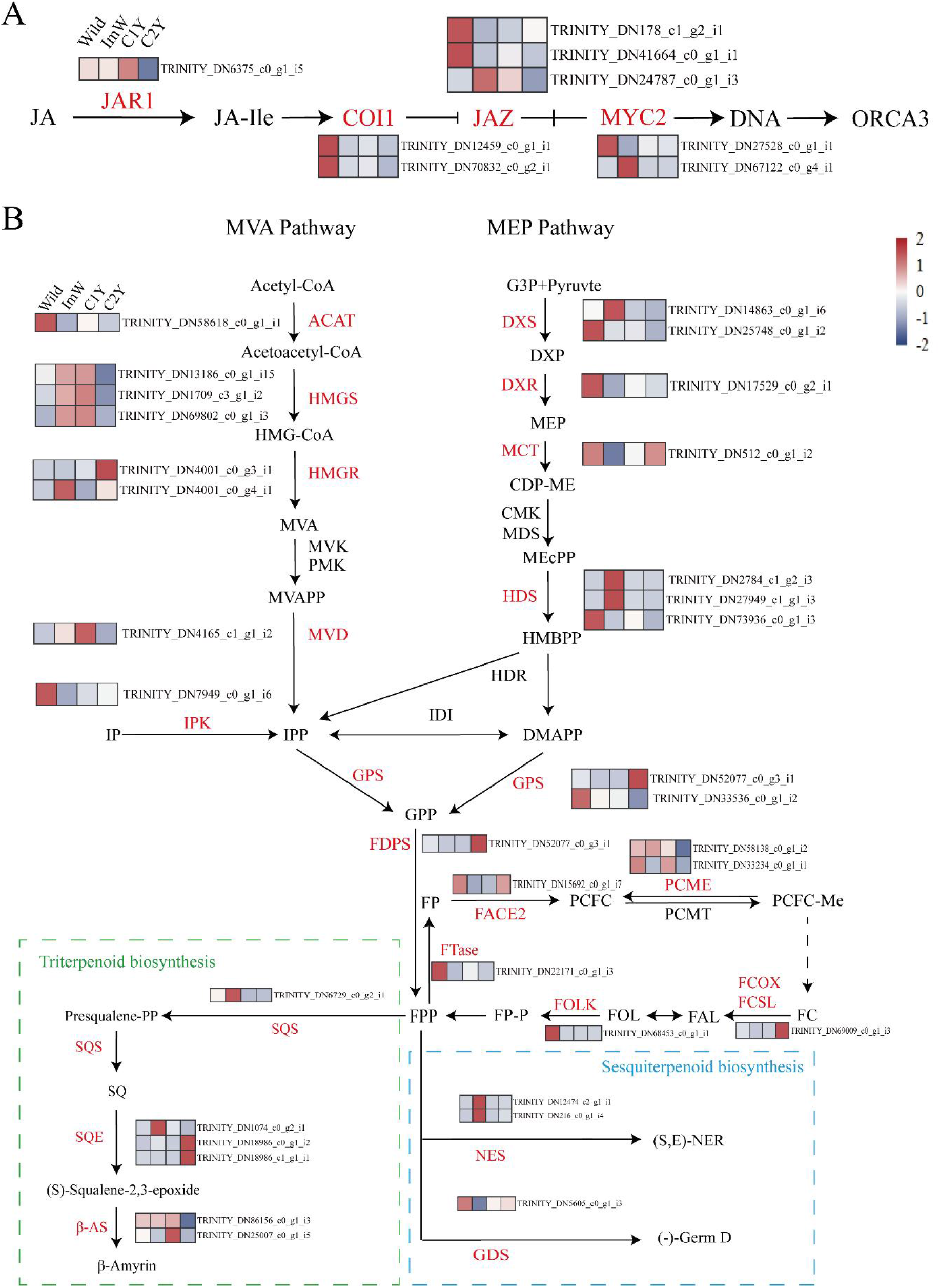
Expression profiles of differentially expressed genes. The FPKM values of the genes were normalized. (A) Expression heatmap of core genes in the JA pathway; (B) Expression patterns of key enzyme genes participating in the MEP, MVA, sesquiterpenoid and triterpenoid biosynthesis pathways. Colour gradient from blue to red corresponded to normalized gene expression values from −2 to 2.

The biosynthesis of sesquiterpenoids in AMR primarily proceeds through the MVA pathway and the MEP pathway, both of which generate the essential intermediate products IPP and DMAPP. Under the action of geranyl diphosphate synthase (GPS) and farnesyl diphosphate synthase (FDPS), IPP and DMAPP could be converted into farnesyl pyrophosphate (FPP), a precursor for sesquiterpenoid synthesis (Sun et al., 2026). In this study, differential expression analysis (threshold: adjusted p-value of FPKM ≤ 0.05) on the groups of C2Y vs Wild, C2Y vs ImW, C2Y vs C1Y and ImW vs Wild identified 33 differentially expressed genes encoding 22 enzymes associated with terpenoid biosynthesis (Fig. 3B) (Table S6). Genes of 1-deoxy-D-xylulose 5-phosphate synthase (DXS), 1-deoxy-D-xylulose 5-phosphate reductoisomerase (DXR), 2-C-methyl-D-erythritol 4-phosphate cytidylyltransferase (MCT) and (E)-4-hydroxy-3-methylbut-2-enyl diphosphate synthase (HDS), which were involved in IPP/DMAPP synthesis in the MEP pathway, exhibited higher expression levels in the Wild group, suggesting that the MEP pathway might be more active in wild AMR. Since pyruvate serves as an upstream carbon-skeleton substrate for the MEP pathway, elevated metabolic flux within the MEP pathway may accelerate pyruvate consumption, which might explain the reduced pyruvate levels observed in wild AMR. (Table S2). In the MVA pathway, the gene encoding acetyl-CoA acetyltransferase (ACAT) was significantly up-regulated in the wild group. This indicated that the catalytic reaction of acetyl-CoA to acetoacetyl-CoA might be more efficient in wild AMR, thereby providing more abundant precursors for the subsequent biosynthesis of IPP.

FPP could undergo cyclization catalyzed by sesquiterpene synthases to form a variety of sesquiterpenoids, and could be converted into various triterpenoids under the action of squalene synthase (SQS), squalene epoxidase (SQE), triterpene cyclases and other enzymes (Guo et al., 2020). Notably, the gene encoding germacrene D synthase (GDS) exhibited the highest expression level in wild AMR, while the gene encoding nerolidol synthase (NES) was expressed at a low level in wild AMR (Fig. 3B). Meanwhile, the genes encoding SQS, SQE, and β-Amyrin synthase (β-AS) were all lowly expressed in the Wild group, implying that more FPP in wild AMR might flow into the sesquiterpenoid synthesis pathway to produce sesquiterpenoids.

Transcription factors (TFs) were screened from transcriptome data. The AP2/ERF-ERF, C2H2 and bHLH families dominated the identified TFs (Fig. S3A). A correlation network was constructed to characterize associations between DEGs participating in sesquiterpenoid biosynthesis and their upstream transcription factors. This network revealed that individual genes were regulated by several upstream transcription factors. (Fig. S3B). DXR, MCT, PCME, DXS, HMGR, HMGS and ACAT corresponded to the larger number of TFs. Among them, DXR was regulated by 39 TFs including MYB, ERF, bHLH and others. MCT was regulated by 36 TFs including MYB, ERF, AUX/IAA and others. PCME was regulated by 37 TFs including PHD, WRKY and others. DXS was regulated by 13 TFs including C2H2, TRAF and others. HMGR was modulated by 19 TFs such as AP2/ERF-ERF and AUX/IAA. A total of 37 TFs (e.g., bHLH and AP2/ERF-ERF) regulated HMGS, whereas ACAT was modulated by 10 TFs including PHD and AP2/ERF-ERF.

### 3.3 Combined analysis of metabolome and transcriptome datasets

To elucidate the associations between gene expression and metabolites and screen core regulatory genes involved in sesquiterpenoid biosynthesis, WGCNA was applied to detect co-expressed gene modules and recognize hub genes. The outcomes were presented in Fig. 4. After filtering out low-expression genes (FPKM < 10), 5953 DEGs were grouped into twelve co-expression modules, and each network cluster was marked by a distinct color (Fig. 4A).

**Fig. 4.**
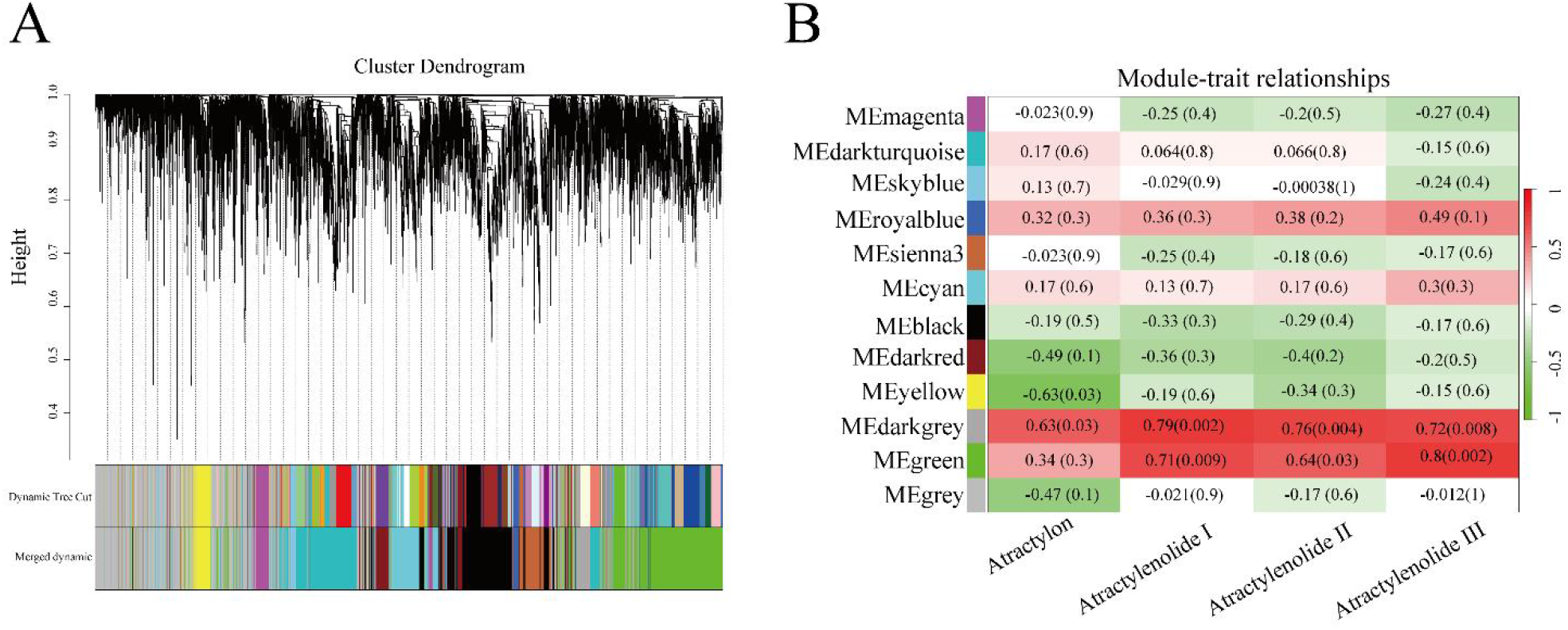
WGCNA clustering module analysis. (A) Co-expression gene modules. (B) Correlation analysis between coexpression gene modules and metabolism traits.

Subsequently, Spearman correlation analysis was performed between co-expression modules and differentially abundant metabolites to screen modules associated with sesquiterpenoid biosynthesis. As shown in Fig. 4B, the results of WGCNA showed that the darkgrey module significantly correlated with the contents of Atractylone and Atractylenolide (Ⅰ, Ⅱ, Ⅲ); nine transcription factors were identified from 203 genes (Table S7), and the hub gene of this module was TRINITY_DN299_c0_g3_i1 (protein transport protein SEC31).

### 3.4 Soil physicochemical properties

Soil nutrients were important for plant growth and development. Soil physicochemical analyses revealed pronounced environmental differentiation among the four groups (Table S8). The rhizosphere soil pH differed significantly among the different groups of AMR, with the imitated wild group exhibiting significantly higher acidity (*P* < 0.05) (Fig. S4A). The soil organic matter content was highest in the wild group samples, followed by the imitated wild group and two-year-old artificially cultivated AMR group, and lowest in one-year-old artificially cultivated AMR group (Fig. S4B). Elemental analysis showed that soil total nitrogen, hydrolyzable nitrogen and available phosphorus contents varied significantly among varieties (*P* < 0.05). The rhizosphere soil of the wild group contained higher total nitrogen, hydrolyzable nitrogen and available phosphorus than the other groups (Fig. S4C-E). These data illustrated that the soil of the wild group was rich in macro-elements.

Mantel’s test was applied to assess associations between soil properties, sesquiterpenoid metabolites, and transcript levels of sesquiterpenoid biosynthetic genes. No significant associations were detected between sesquiterpenoid metabolites and macronutrients (N and P), soil pH, or soil organic matter. In contrast, only pH was correlated with the expression of sesquiterpenoid biosynthetic genes (Fig. S5). These observations indicated that such edaphic variables were not major contributors to sesquiterpenoid accumulation.

### 3.5 Analysis of endophytic and rhizospheric microorganisms in different varieties of AMR

To further elucidate the regulation of differential sesquiterpenoid metabolites by AMR root-associated bacterial and fungal communities, sequencing and taxonomic annotation were conducted on microbial assemblages recovered from root and rhizosphere soil samples of AMR grown under different growth environments. The results of alpha diversity analysis on endophytic and rhizosphere bacteria and fungi among these groups were shown in Fig. S6. The Chao1 and Observed species indices were primarily applied to evaluate species richness. The Shannon and Simpson indices mainly reflected species diversity. For endophytic bacteria of AMR, the Shannon, Simpson, Chao1 and Observed species indices of the imitative wild AMR were higher than those of the other groups, with significant differences compared with one-year artificially cultivated AMR (*P* < 0.05) (Fig. S6A). For rhizosphere bacteria of AMR, the Shannon, Simpson, Chao1 and Observed species indices of the wild group were higher than those of all cultivated groups. However, no significant differences in the Chao1 and Observed species indices were detected between the wild group and other groups, while the Shannon and Simpson indices showed significant differences relative to one-year artificially cultivated AMR (*P* < 0.05), which suggested that the wild AMR harbored higher species diversity of rhizosphere bacteria (Fig. S6B). For endophytic fungi, the one-year-old artificially cultivated group exhibited higher Shannon, Simpson, Chao1 and Observed species indices relative to the remaining groups. The Chao1 and Observed species indices exhibited significant differences compared with wild and imitative wild AMR (*P* < 0.05), and the Shannon and Simpson indices differed significantly from imitative wild and two-year artificially cultivated AMR (*P* < 0.05) (Fig. S6C). For rhizosphere fungi, significant differences in the Chao1 and Observed species indices were observed between imitative wild and artificially cultivated AMR (*P* < 0.05), while no marked intergroup variations were detected for the Shannon and Simpson indices (Fig. S6D). Notably, rhizosphere bacteria exhibited consistently greater species richness than endophytic bacteria in all tested samples.

The results of PCoA based on Bray-Curtis distance was showed in Fig. S6. The results revealed that the rhizosphere bacterial communities from distinct AMR types exhibited stronger separation (R² = 0.64, *P* = 0.001) (Fig. S7B) compared with endophytic bacterial communities (R² = 0.44, *P* = 0.028) (Fig. S7A). Similarly, the separation of rhizosphere fungal communities (R² = 0.56, *P* = 0.001) (Fig. S7D) was significantly higher than that of endophytic fungal communities (R² = 0.36, *P* = 0.004) (Fig. S7C).

Analysis of the top 10 root-associated bacterial and fungal species in AMR at the phylum and genus levels revealed that endophytic bacteria from the four AMR groups were predominantly classified within *Proteobacteria*, *Actinobacteriota*, and *Firmicutes*. *Firmicutes* showed greater relative abundance within the wild group relative to the remaining three groups. *Actinobacteriota* dominated the two-year-old artificially cultivated group among all groups, and the one-year-old artificially cultivated group exhibited the highest relative abundance of *Proteobacteria* among all groups (Fig. 5A). *Actinobacteriota*, *Proteobacteria*, and *Acidobacteriota* were the dominant phyla associated with the rhizosphere bacterial community. *Acidobacteriota* attained the greatest relative abundance within the wild group across all four groups. *Proteobacteria* were most enriched in the imitative wild group, whereas *Actinobacteriota* were most enriched in the two-year artificially cultivated group. (Fig. 5B). Notably, the two communities exhibited significant similarity in phylum-level composition. On the genus taxonomic level, the dominant taxa within endophytic bacterial communities were *Delftia*, which exhibited the highest relative abundance in one-year-old artificially cultivated AMR, while the genus *Burkholderia-Caballeronia-Paraburkholderia* was the most abundant in two-year-old artificially cultivated AMR (Fig. 5C). Rhizosphere bacterial communities showed no single dominant genus, with distinct differences between the two communities. *Vicinamibacteraceae* were most abundant in the wild group, *Acidothermus* was the most abundant in the imitative wild group, *Pseudarthrobacteraceae* exhibited the highest relative abundance in the one-year-old artificially cultivated group, and *Bacillus* had the highest relative abundance in the two-year-old artificially cultivated group (Fig. 5D).

**Fig. 5.**
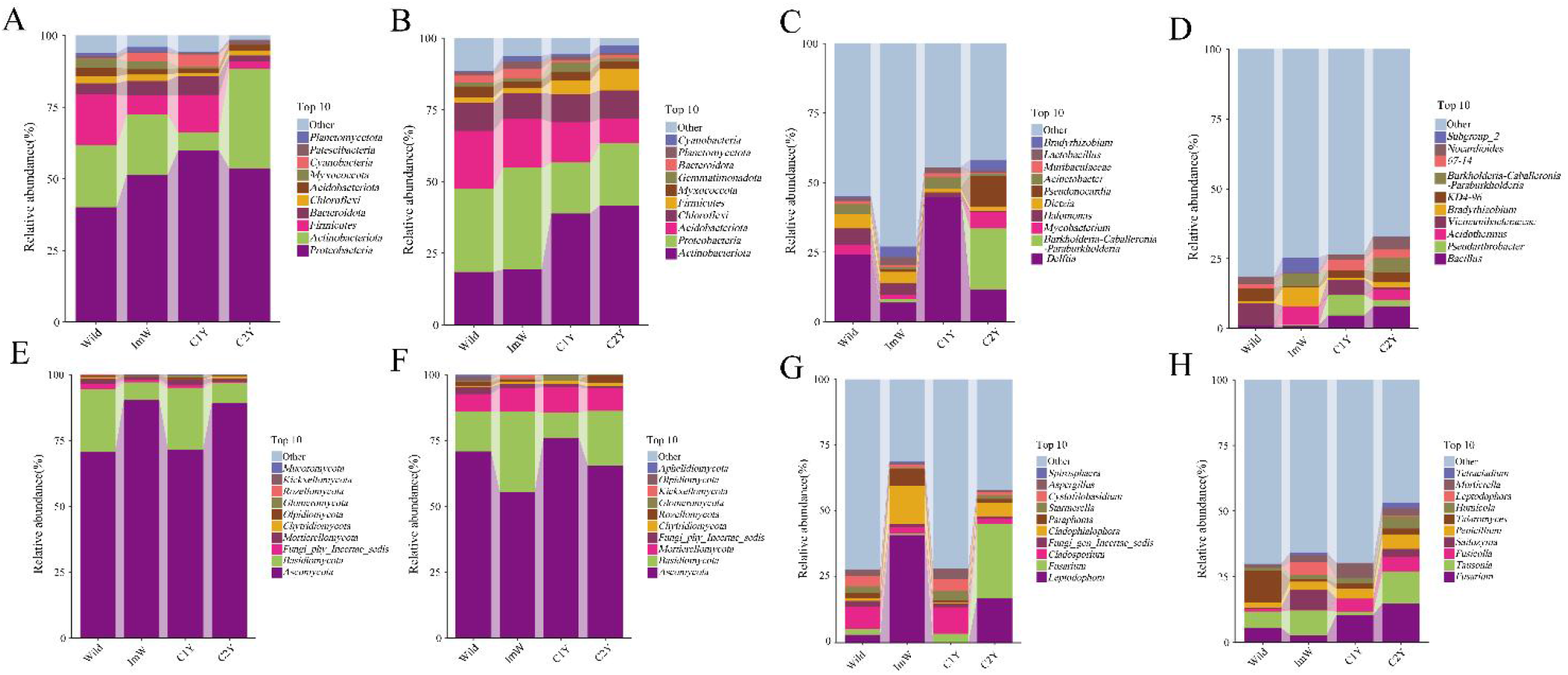
Community composition of root-associated bacteria and fungi in AMR under different cultivation modes. (A – B) Endophytic (A) and rhizosphere (B) bacterial phylum-level communities; (C – D) endophytic (C) and rhizosphere (D) bacterial genus-level communities; (E–F) endophytic (E) and rhizosphere (F) fungal communities at the phylum level; (G–H) endophytic (G) and rhizosphere (H) fungal communities at the genus level. Wild, ImW, C1Y and C2Y represented wild AMR, imitative wild-cultivated AMR, one-year artificially cultivated AMR, and two-year artificially cultivated AMR, respectively. The number of biological replicates was three.

At the phylum level, for endophytic fungi, *Ascomycota* and *Basidiomycota* dominated the fungal communities. *Ascomycota* were most abundant in the imitative wild group, and *Basidiomycota* showed the highest relative abundance in the wild group versus the other three groups (Fig. 5E). *Ascomycota*, *Basidiomycota*, and *Mortierellomycota* dominated the rhizosphere fungal communities. *Ascomycota* exhibited the highest relative abundance in the one-year-old artificially cultivated group. *Basidiomycota* showed greater relative abundance in the imitative wild group than in the other groups, and *Mortierellomycota* was more abundant in the one-year-old artificially cultivated group than in the remaining three groups (Fig. 5F). The endophytic fungal communities and the rhizosphere fungal communities exhibited distinct differences in genus-level species composition. For the endophytic fungal communities, the relative abundances of *Leptodophora* and *Cladophialophora* in the imitative wild group were higher than those in the other three groups, and *Fusarium* was more abundant in the two-year-old artificially cultivated group than in the other three groups (Fig. 5G). For rhizosphere fungal communities, the two-year-old artificially cultivated group had higher relative abundances of *Fusarium* and *Tausonia* than the other three groups, while *Talaromyces* was most abundant in the wild group. (Fig. 5H).

LEfSe analysis of the endophytic and rhizosphere bacteria and fungi of AMR revealed 6, 29, 12, and 39 specific marker genera for endophytic bacteria, rhizosphere bacteria, endophytic fungi, and rhizosphere fungi, respectively, suggesting marked divergence in the recruitment patterns of bacterial and fungal communities across distinct AMR groups. Notably, endophytic bacteria exhibited the most pronounced enrichment in two-year-old cultivated AMR, while no significant enrichment was detected in one-year-old cultivated and wild AMR. In contrast, wild AMR harbored higher abundances of rhizosphere bacteria. For endophytic bacteria, *Obscuribacteraceae* served as the dominant marker taxon in the imitative wild group, while *Burkholderia-Caballeronia-Paraburkholderia* was the most representative marker genus in the two-year-old artificially cultivated group (Fig. S8A). Among rhizosphere bacteria, *Nitrospira*, *Bradyrhizobium*, *Pseudarthrobacter* and *Bacillus* were the dominant marker genera for the wild, imitative wild, one-year-old artificially cultivated and two-year-old artificially cultivated groups, respectively (Fig. S8B). For endophytic fungi, the characteristic marker genera were *Debaryomyces*, *Cladosporium* and *Fusarium* in the wild, one-year-old artificially cultivated and two-year-old artificially cultivated groups, correspondingly (Fig. S8C). As for rhizosphere fungi, *Talaromyces*, *Saitozyma*, *Alternaria* and *Fusarium* were identified as the dominant marker genera in the wild, imitative wild, one-year-old artificially cultivated and two-year-old artificially cultivated groups, respectively (Fig. S8D).

### 3.6 The effect of soil physicochemical properties and sesquiterpenoid metabolites on the root microbial communities

The correlations of soil properties and sesquiterpenoid metabolites with the microbial community structure were performed by the Mantel test. As shown in Fig. 6, endophytic and rhizosphere microbiota exhibited distinct response patterns to soil physicochemical properties and differential sesquiterpene metabolites. Endophytic bacteria and endophytic fungi were only significantly correlated with soil pH and available phosphorus (*P* < 0.05). Rhizosphere bacteria were significantly associated with pH, hydrolyzable nitrogen and available phosphorus (*P* <0.05). The rhizosphere fungi were significantly correlated with pH, hydrolyzable nitrogen, total nitrogen, organic matter and available phosphorus (*P* <0.05), and rhizosphere fungi were most sensitive to environmental changes. Neither endophytic bacteria nor fungi exhibited significant correlations with these differential sesquiterpenes. Rhizosphere bacteria were significantly correlated with atractylon, and rhizosphere fungi showed significant correlations with valencene, (s)-β-bisabolene, atractylon, and atractylenolide (I, II, III) (*P* <0.05). Collectively, soil factors and sesquiterpenoid metabolites could impact the microbial composition of rhizosphere soil, while the microbial composition of rhizosphere fungi were more associated with the soil physicochemical properties and sesquiterpenoid metabolites.

**Fig. 6.**
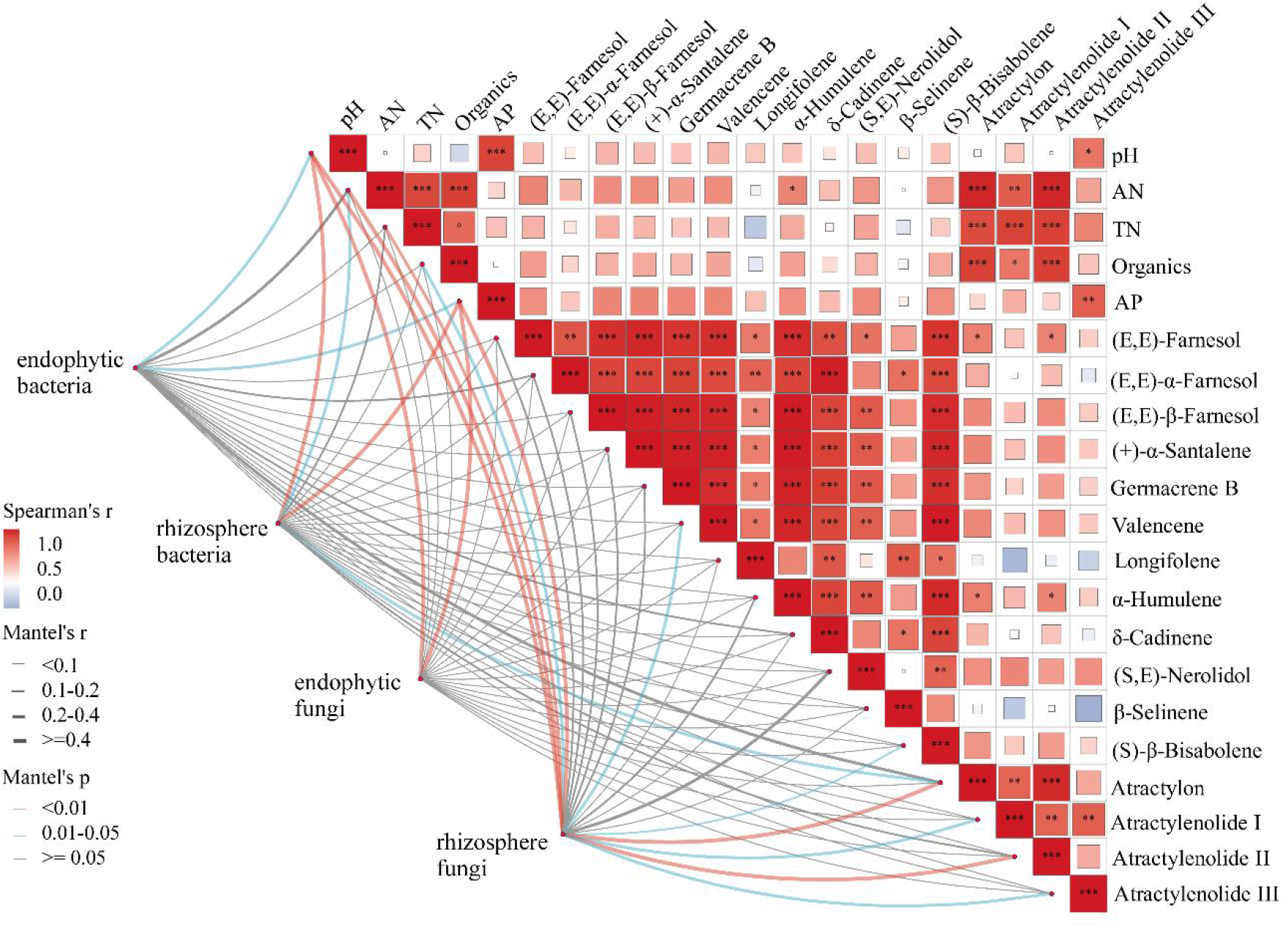
Visualization of correlation heatmap and Mantel test revealing linkages of rhizosphere and endospheric microbial communities with edaphic properties and metabolite abundance patterns. Color gradients in the heatmap indicate Pearson correlation coefficients; line width and color separately reflect Mantel’s r and Mantel’s *p* values.

### 3.7 Multi-omics correlation analysis

To further explore the interactions among rhizosphere microorganisms, sesquiterpenoid-related differentially expressed genes, and differential sesquiterpenoid metabolites in AMR, we conducted a correlation analysis involving the top 20 abundant bacterial and fungal taxa from the endosphere and rhizosphere, sesquiterpenoid-related differentially expressed genes, and differential sesquiterpenoid metabolites. Correlation coefficients linking bacterial relative abundance, metabolite concentrations and gene expression profiles were computed via Spearman correlation analysis. *P* values were calculated to evaluate statistical significance, with Benjamini–Hochberg correction applied for multiple comparisons to limit the false discovery rate. The outputs of correlation analysis were visualized in Figure 7. Figure 7 displayed an interdomain association network integrating microbiome, transcriptome and metabolomic datasets. This network was built upon the above correlation outputs; nodes corresponded to bacterial genera, metabolites or genes, whereas edges denoted statistically significant associations (*P* < 0.05 and FDR < 0.05 after Benjamini–Hochberg adjustment) (Table S9). For endophytic bacteria, atractylone and atractylenolide (Ⅰ, Ⅱ, Ⅲ), as well as the DXS-coding gene TRINITY_DN25748_c0_g1_i2, showed a marked positive association with the abundance of *Pseudomonas* (*P* < 0.05). Similarly, atractylone and atractylenolide (Ⅰ, Ⅲ) exhibited a marked positive association with the relative abundance of *Halomonas* (*P* < 0.05). The contents of atractylenolid Ⅲ, together with the genes encoding enzymes including HMGS (TRINITY_DN13186_c0_g1_i15), HDS (TRINITY_DN73936_c0_g1_i3), PCME (TRINITY_DN33234_c0_g1_i1), and FTase (TRINITY_DN22171_c0_g1_i3), showed significantly positive correlations with *Acinetobacter* (*P* < 0.05). For endophytic fungi, atractylone and atractylenolide (Ⅰ, Ⅱ, Ⅲ), as well as DXS (TRINITY_DN25748_c0_g1_i2) and HDS (TRINITY_DN73936_c0_g1_i3) showed significantly positive correlations with the relative abundance of *Fungi_gen_Incertae_sedis* (*P* < 0.05). The enzyme-coding gene PCME (TRINITY_DN33234_c0_g1_i1) and HMGS (TRINITY_DN13186_c0_g1_i15), as well as atractylone exhibited a marked positive association with the relative abundance of *Aspergillus* (*P* < 0.05). For rhizosphere bacteria, the contents of atractylone and atractylenolide II, together with the genes encoding enzymes including HMGR (TRINITY_DN4001_c0_g4_i1) and NES (TRINITY_DN12474_c2_g1_i1) exhibited a marked positive association with the relative abundance of *IMCC26256* (*P* < 0.05). The contents of atractylenolid Ⅲ, together with the genes encoding enzymes, including HMGR (TRINITY_DN4001_c0_g4_i1), HDS (TRINITY_DN2784_c1_g2_i3) and NES (TRINITY_DN12474_c2_g1_i1 and TRINITY_DN216_c0_g1_i4) exhibited a marked positive association with the relative abundance of *Subgroup2* (*P* < 0.05). *Vicinamibacteraceae* was significantly positively correlated with atractylenolide ⅡI and multiple enzyme-encoding genes, including DXS (TRINITY_DN25748_c0_g1_i2), HDS (TRINITY_DN73936_c0_g1_i3), PCME (TRINITY_DN33234_c0_g1_i1), FOLK (TRINITY_DN68453_c0_g1_i1), ACAT (TRINITY_DN58618_c0_g1_i1), DXR(TRINITY_DN17529_c0_g2_i1), IPK (TRINITY_DN7949_c0_g1_i6), FTase (TRINITY_DN22171_c0_g1_i3) and GDS (TRINITY_DN5605_c0_g1_i3) (*P* < 0.05). *Rokubacterides* was significantly positively correlated with atractylenolide (I, ⅡI) and multiple enzyme-encoding genes, including DXS (TRINITY_DN14863_c0_g1_i6 and TRINITY_DN17529_c0_g2_i1), HMGS (TRINITY_DN13186_c0_g1_i15), PCME (TRINITY_DN33234_c0_g1_i1), HDS (TRINITY_DN73936_c0_g1_i3), ACAT (TRINITY_DN58618_c0_g1_i1) and FTase (TRINITY_DN22171_c0_g1_i3) (*P* < 0.05). *Acidibacter* was correlated positively and markedly with atractylone and multiple enzyme-encoding genes, including HMGR (TRINITY_DN4001_c0_g4_i1) and NES (TRINITY_DN12474_c2_g1_i1 and TRINITY_DN216_c0_g1_i4) (*P* < 0.05). For rhizosphere fungi, atractylone and atractylenolide II, as well as HMGR (TRINITY_DN4001_c0_g4_i1) showed significantly positive correlations with the relative abundance of *Thelonectria* (*P* < 0.05). Atractylenolide (I, II), DXS (TRINITY_DN14863_c0_g1_i6) and PCME (TRINITY_DN33234_c0_g1_i1 and TRINITY_DN58138_c0_g1_i2) showed significantly positive correlations with the relative abundance of *Rhizoctonia* (*P* < 0.05). Atractylone and atractylenolide (Ⅰ, Ⅱ, Ⅲ), as well as DXS (TRINITY_DN25748_c0_g1_i2) and FOLK (TRINITY_DN68453_c0_g1_i1) showed significantly positive correlations with the relative abundance of *Fungi_gen_Incertae_sedis* (*P* < 0.05). *Aspergillus* was correlated positively and markedly with germacrene B and multiple enzyme-encoding genes, including FACE2 (TRINITY_DN15692_c0_g1_i7), MCT (TRINITY_DN512_c0_g1_i2), ACAT (TRINITY_DN58618_c0_g1_i1), DXR(TRINITY_DN17529_c0_g2_i1), IPK (TRINITY_DN7949_c0_g1_i6), FTase (TRINITY_DN22171_c0_g1_i3) and GDS (TRINITY_DN5605_c0_g1_i3) (*P* < 0.05). *Talaromyces* was significantly positively correlated with germacrene B, atractylenolide ⅡI and multiple enzyme-encoding genes, including FACE2 (TRINITY_DN15692_c0_g1_i7), PCME (TRINITY_DN33234_c0_g1_i1), FOLK (TRINITY_DN68453_c0_g1_i1), ACAT (TRINITY_DN58618_c0_g1_i1), DXR(TRINITY_DN17529_c0_g2_i1), IPK (TRINITY_DN7949_c0_g1_i6), MCT (TRINITY_DN512_c0_g1_i2) and GDS (TRINITY_DN5605_c0_g1_i3) (*P* < 0.05).

**Figure 7.**
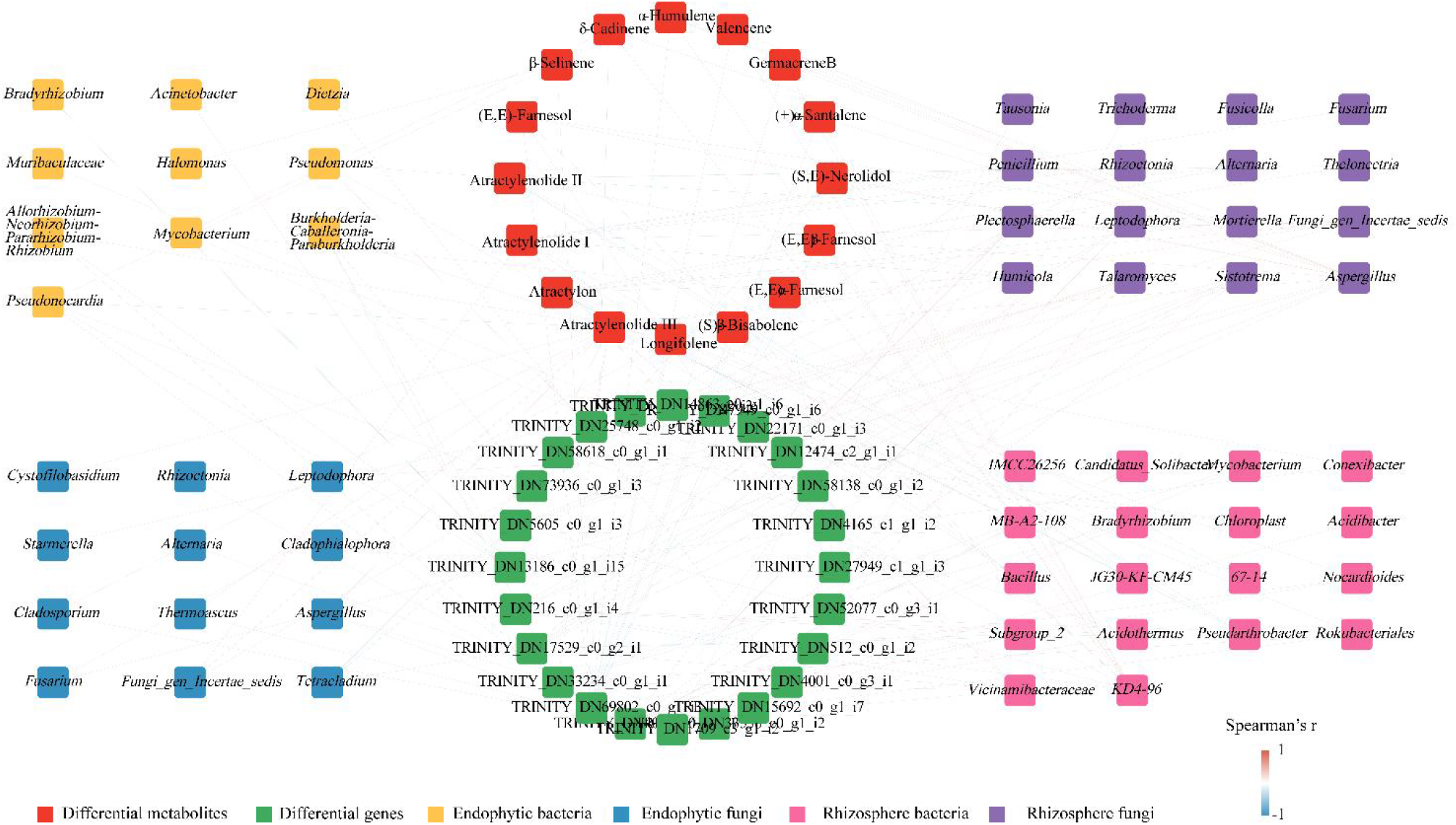
Spearman correlation network among differential metabolites, differentially expressed genes, endophytic microbes and rhizosphere microbes of AMR. Nodes represent differential metabolites (red), differentially expressed genes (green), endophytic bacteria (yellow), endophytic fungi (blue), rhizosphere bacteria (pink), and rhizosphere fungi (purple). The color gradient of connecting lines indicates the value of Spearman’s correlation coefficient (r), ranging from −1 to 1.

Meanwhile, multiple enzyme-encoding genes including DXS (TRINITY_DN14863_c0_g1_i6), HDS (TRINITY_DN73936_c0_g1_i3), FOLK (TRINITY_DN68453_c0_g1_i1) and PCME (TRINITY_DN58138_c0_g1_i2) showed significantly positive correlations with atractylenolide I (*P* < 0.05). Multiple enzyme-encoding genes including DXS (TRINITY_DN25748_c0_g1_i2), HMGS (TRINITY_DN13186_c0_g1_i15), HDS (TRINITY_DN73936_c0_g1_i3), PCME (TRINITY_DN33234_c0_g1_i1),DXR(TRINITY_DN17529_c0_g2_i1) and FTase (TRINITY_DN22171_c0_g1_i3) showed significantly positive correlations with atractylenolide III (*P* < 0.05). Germacrene B showed significantly positive correlations with multiple enzyme-encoding genes including IPK ( TRINITY_DN7949_c0_g1_i6), MCT (TRINITY_DN512_c0_g1_i2) and GDS (TRINITY_DN5605_c0_g1_i3) (*P* < 0.05).

## Discussion

This work adopted an integrated analysis of widely targeted metabolomics, transcriptome and rhizosphere microbiome to systematically explore how growth years and cultivation patterns (wild, imitated wild, and artificially cultivated) affect medicinal component accumulation, gene expression, and root-associated microbial communities of AMR. The results showed that AMR from different sources exhibited significant differentiation at the metabolomic, transcriptomic, and microbiomic levels, and there was a close regulatory correlation among these three aspects.

Sesquiterpenoids, especially atractylone and atractylenolides (Ⅰ, Ⅱ and Ⅲ), were the most important bioactive components of AMR. However, their compositions and differences in different growth years and cultivation conditions remained poorly characterized. In this study, metabolic quantification revealed that wild AMR accumulated significantly higher levels of these sesquiterpenoids than imitated wild and artificially cultivated counterparts. In addition, two-year-old cultivated AMR exhibited markedly higher atractylone abundance than one-year-old plants, which was consistent with previously documented trends (Song et al., 2023; Zhao et al., 2021).

Primary metabolism supplies carbon precursors for sesquiterpenoid biosynthesis through the MVA and MEP pathways (Lee et al., 2016; Rohmer, 1999). Untargeted metabolomic data showed that wild AMR may consume more pyruvate for sesquiterpenoid synthesis, and this finding was corroborated by transcriptomic results. Based on these untargeted metabolomic findings, wild and imitated-wild AMR could redirect primary carbon metabolism to supply sufficient precursors for the MEP and MVA pathways, yielding higher total sesquiterpenoid accumulation than two-year artificially cultivated AMR. The elevated fructose-6-phosphate level observed in one-year-old cultivated AMR may relate to plant growth status.

In the widely-targeted metabolomics analysis, the differential metabolites were significantly enriched in the sesquiterpenoid and triterpenoid biosynthetic pathways. And it had revealed that the pathway was the core route determining the medicinal quality of AMR during the biosynthesis of sesquiterpenoids (Cui et al., 2025). Notably, in the ImW vs Wild group, the result of the widely targeted metabolomics analysis revealed that the content of germacrene B in wild AMR was significantly higher than that in imitated wild AMR, and also higher than that in artificially cultivated AMR. It had found that the content of atractylon in *Atractylodes lancea* was consistent with the variation in germacrene B content, leading to the speculation that germacrene B was a precursor substance for atractylon synthesis (Zhu et al., 2025). Combined with our metabolomic data, the higher level of germacrene B detected in wild AMR might facilitate massive accumulation of atractylone in wild AMR. The distinct separation of metabolic profiles among the four AMR groups suggested that growth years and cultivation age could reshape the overall metabolic flux and drive more carbon resources toward secondary metabolism. The significant accumulation of atractylone, atractylenolide Ⅰ, and atractylenolide Ⅱ in wild AMR might be attributed to the preferential allocation of metabolic flux to sesquiterpenoid biosynthesis under natural habitats, which also represented the key reason for the superior quality of authentic medicinal materials.

At the transcriptional regulation level, differentially expressed genes exhibited significant enrichment in pathways including plant hormone signal transduction, plant MAPK signaling pathway, and photosynthesis. Combined with the metabolism results, the accumulation of sesquiterpenoids might be influenced by the primary metabolic pathways linked to sesquiterpenoids synthesis. Among the plant hormone signal transduction pathway, the JA signaling pathway acted as a core node regulating the biosynthesis of sesquiterpenoid skeletons (Chen et al., 2019; Yang et al., 2019). The expression levels of COI1, JAZ and MYC2 in the wild group were higher than those in other groups, which might further promote the synthesis of downstream sesquiterpenoids (Zhou & Memelink, 2016). The biosynthesis of sesquiterpenoids in AMR mainly depended on the MVA pathway and the MEP pathway. In the MEP pathway, key genes including DXS, DXR, MCT and HDS, which governed the biosynthesis of IPP/DMAPP precursors, were highly expressed in wild samples. The results indicated that the MEP pathway was more active in wild AMR and could provide sufficient precursors for sesquiterpenoid synthesis (Park et al., 2022). By contrast, the genes SQS, SQE and β-AS were expressed at low levels in the wild group, which reduced the carbon flux allocated to the triterpenoid biosynthesis branch (Mazzuca et al., 2014). Consequently, more FPP was diverted into the sesquiterpenoid synthesis pathway, ultimately facilitating the efficient accumulation of bioactive ingredients. WGCNA identified a darkgrey module significantly correlated with the contents of atractylenolide and atractylone, and the hub gene of this module was TRINITY_DN299_c0_g3_i1, which encoded the protein transport protein Sec31. Sec31 was a core component of the COPII coat complex and played an essential role in ER-to-Golgi protein trafficking (Nie & Ma, 2025). Moreover, it had reported that the key biosynthetic and modifying enzymes for sesquiterpenoids, such as HMGR and cytochrome P450 enzymes, were all distributed in the endoplasmic reticulum membrane (Erffelinck & Goossens, 2018). Therefore, high expression of SEC31 might indirectly influence the biosynthetic efficiency of sesquiterpenoids.The sesquiterpenoids contents of medicinal plants could be impacted by soil conditions and the plant-associated microbiome. Soil nutrients could impact the bioactive components of medicinal plants (Lim et al., 2012; Stewart et al., 2002) and influence the soil microbial composition (Jiang et al., 2017; Jiao et al., 2018). The relationship between soil properties with sesquiterpenoid metabolites and the expression of genes related to its biosynthesis showed that these environmental parameters (soil pH, organic matter, nitrogen, and phosphorus) might not be significant factors in enhancing the production of sesquiterpenoid. The significant differences in soil pH, organic matter, nitrogen, and phosphorus among the four AMR groups indicated that distinct soil environments under different cultivation modes could give rise to variations in the microbial communities of AMR, thereby affecting the accumulation of its medicinal active components. The result was consistented with the previous study that soil conditions could shape the rhizosphere microbial communities, which in turn regulate the biosynthesis and accumulation of secondary metabolites in medicinal plants (Wang et al., 2022).

Consistent with general plant microbiome patterns (Beckers et al., 2017; Ha et al., 2021; Li et al., 2026), rhizosphere bacterial diversity exceeded endophytic bacterial diversity in AMR, consistent with the host filtering model of root microbiome assembly (Trivedi et al., 2020). At the genus level, rhizosphere and endophytic bacterial and fungal biomarkers differed markedly among AMR cultivars. Meanwhile, the community compositions of the top 10 most abundant bacterial and fungal genera also displayed marked intergroup differences. Mounting studies had confirmed that plant environment background and metabolic characteristics could reshape root-associated microbiomes, and in turn, microbes could modulate plant secondary metabolite biosynthesis (Liu et al., 2022; Rodríguez et al., 2025). The relationships between soil properties and sesquiterpenoid metabolites with the microbial community structure were performed by Mantel test. The results showed that soil factors and sesquiterpenoid metabolites could impact the microbial composition of rhizosphere soil, while the microbial composition of rhizosphere fungi was more associated with the soil physicochemical properties and sesquiterpenoid metabolites. Consistent with our findings, prior work had demonstrated that rhizosphere fungi were generally more sensitive to soil environmental changes than bacteria (Xu et al., 2019). Root-derived terpenoids are known to act as selective chemical signals that recruit specific rhizosphere microorganisms, with fungal communities generally more responsive than bacterial communities (Carvalhais et al., 2019; Pang et al., 2021). Although root-exuded sesquiterpenoids were not directly measured in this study, the significant correlations between root sesquiterpenoid accumulation and the abundances of specific rhizosphere taxa lead us to further hypothesize that partial root-synthesized sesquiterpenoids or their metabolic intermediates might be released into the rhizosphere via root exudation and serve as cross-compartment signals participating in the regulation of rhizosphere microbiota assembly.

Accumulating evidence recognized the critical role of microbes in modulating plant secondary metabolism (Qin et al., 2022; Yanmei et al., 2023). For example, microbes could enhance the content of furanocoumarins in *Angelica dahurica* var. *Formosana* (Jiang et al., 2024), and endophytic fungi in *Angelica sinensis* could induce the production of novel metabolites (Chen et al., 2025). In this study, we performed an integrated analysis of the metabolome, transcriptome, and microbiome to reveal a close correlation between the root microbiota and sesquiterpenoid accumulation in AMR. The results showed that the relative proportion of multiple microbial taxa, including endophytic bacteria (*Pseudomonas*, *Halomonas*, *Acinetobacter*), endophytic fungi (*Fungi_gen_Incertae_sedis*, *Aspergillus*), rhizosphere bacteria (*IMCC26256*, *Subgroup 2*, *Acidibacter*, *Rokubacteriales*, *Vicinamibacteraceae*), and rhizosphere fungi (*Aspergillus*, *Talaromyces*, *Fungi_gen_Incertae_sedis*, *Thelonectria*, *Rhizoctonia*), were significantly and positively correlated with the contents of core sesquiterpenoids (such as atractylone and atractylenolides) and the expression levels of key terpenoid biosynthetic genes (e.g., HMGS, HDS, PCME, DXS, ACAT, FTase, HMGR) (*P* < 0.05). These correlations suggested that microbial partners not only coexisted with the host but were metabolically intertwined with its secondary metabolite regulation. Previous studies had reported that *Pseudomonas* could promote the accumulation of atractylone and atractylenolides (Yang et al., 2019). Additionally, fungi of the genus *Aspergillus* possessed a complete sesquiterpene biosynthetic pathway and were capable of synthesizing various sesquiterpenoids (Zhao et al., 2022). Functional verification experiments had demonstrated that the endophytic *Acinetobacter* sp. ALEB16 isolated from *Atractylodes lancea* significantly upregulated the key terpenoid biosynthetic gene HMGR and facilitated the biosynthesis of sesquiterpenoids such as atractylone mainly via the MVA pathway (Wang et al., 2014). Consistently, multi-omics correlation analysis in this study revealed a significant positive correlation between *Acinetobacter* and HMGS, a core gene of the MVA pathway, implying that this genus also had regulatory potential for sesquiterpenoid biosynthesis in AMR. Previous rhizosphere microbiome association research on *Atractylodes chinensis* revealed that *Vicinamibacteraceae* and *Thelonectria* exhibited significant positive associations between their relative abundance and the concentrations of sesquiterpenoid active constituents (Bai et al., 2025). This correlation pattern was consistent with our findings regarding rhizosphere microbiota and active compounds of AMR. Accordingly, we speculate that these two microbial taxa might be involved in the accumulation of sesquiterpenoids in AMR, though their underlying regulatory mechanisms remained to be verified. Taken together, we propose a stratified working model: under different cultivation types, soil physicochemical properties are not the primary direct trigger driving sesquiterpenoid biosynthesis in AMR. Instead, rhizosphere microorganisms were the key contributors. Rhizosphere microorganisms might promote sesquiterpenoid accumulation by regulating the expression of key enzyme genes involved in sesquiterpenoid biosynthesis. This process formed a “microbe-gene-sesquiterpenoid” metabolic positive-feedback loop, which collectively supported the accumulation of bioactive sesquiterpenoids in AMR. It should be noted that this stratified model was inferred from multi-omics correlative data, and definitive causal relationships await further functional experimental validation. The differences in the enrichment of key beneficial microbial communities and the expression patterns of terpenoid pathway genes among wild, imitated wild, and artificially cultivated AMR collectively determined the differential accumulation of active components such as atractylone and atractylenolides. Future research should focus on elucidating how these microorganisms further influence plant gene expression and the biosynthesis of secondary metabolites. These findings will provide critical insights for developing strategies to modulate root microbiomes, thereby improving plant performance and boosting the yield of pharmaceutically valuable metabolites.

## Conclusion

This work adopted a multi-omics strategy to explore the intricate crosstalk between sesquiterpenoid secondary metabolites, genes encoding key enzymes for sesquiterpenoid biosynthesis, as well as rhizosphere and endophytic microbiomes of AMR across distinct growth environments. The results demonstrated that the accumulation of sesquiterpenoids, expression of terpenoid biosynthetic genes, and community structure of endophytic and root-associated microorganisms differed significantly in AMR under various growth environments. Correlation analysis demonstrated that the abundances of specific microbial taxa were tightly associated with the contents of core active components, including atractylone and atractylenolides (I, II, III), as well as the expression levels of key sesquiterpenoid biosynthetic genes. Importantly, rhizosphere microorganisms acted as critical regulators driving sesquiterpenoid accumulation in AMR, while sesquiterpenoid metabolites in turn modulated the assembly and composition of rhizosphere microbial communities, forming a bidirectional plant–microbe interaction network. Although this study is based on correlational analyses with preliminary conclusions, the results provide valuable insights into the intricate relationships between AMR and its root-associated microbiome. Further experimental studies are needed to elucidate the precise mechanisms underlying these interactions. Ultimately, understanding the interactions between AMR and its microbiome at the molecular level will help develop optimized cultivation strategies and improve the accumulation of active secondary metabolites in this medicinal plant. The present study thus provides a mechanistic foundation for future efforts to engineer the root microbiome as a strategy to enhance the quality and yield of medicinal plants.

## Acknowledgements

This work was supported by the Key Project of Natural Science Research in Anhui Universities (nos. 2024AH050975, 2024AH050978 and KJ2021A0593), Major Scientific Research Project of Universities in Anhui Province (2025AHGXZK20055), the Joint Research Center for Chinese Herbal Medicine of Anhui of IHM (no. Yjzx2023008), and the Foundation of Anhui Engineering Research Center for Quality Improvement and Utilization of Genuine Chinese Medicinal Materials (no. gcyj202407).

## CRediT authorship contribution statement

**Meili Chen:** Writing – original draft, Supervision, Project administration, Formal analysis. **Shiqi Yu:** Software, Investigation. **Yakun Wang:** Visualization, Investigation. **Yi Zheng:** Validation, Data curation. **Rongxin Zou:** Investigation, Data curation. **Juan Liang:** Investigation, Validation. **Wentao Zhu:** Software, Data curation. **Wei Fang:** Visualization, Validation, Software. **Dianlei Wang:** Investigation, Data curation. **Nianxia Sun:** Writing – review & editing, Resources, Funding acquisition, Conceptualization.

## Data Availability

Data will be made available on request.

