## Supplemental Data 1 for "Multi-omics reveals accumulation mechanisms underlying differences in sesquiterpenoid content across different growth environments of *Atractylodes macrocephala* rhizome"

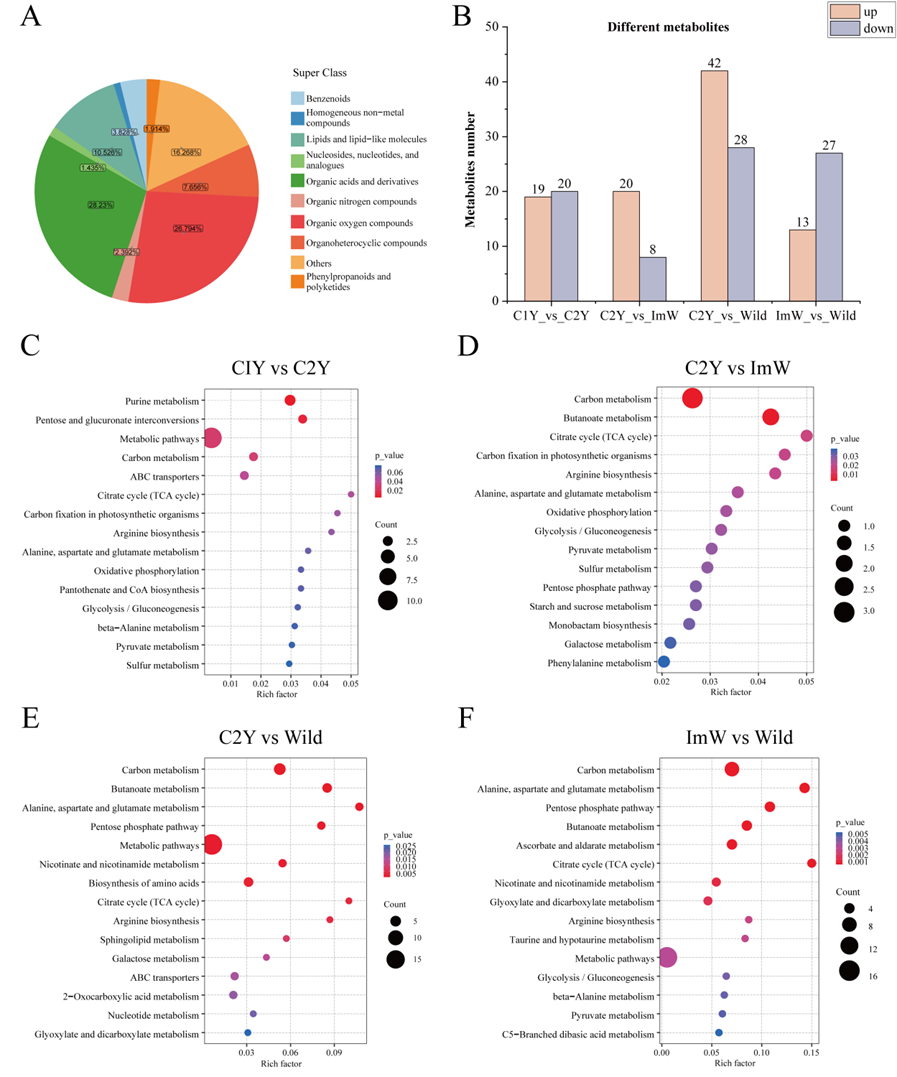

Fig. S1 Metabolomic profiling analysis of key bioactive components in four AMR sample groups via untargeted metabolomics analysis. (A) The classification of all identified metabolites. (B) Differences in metabolite contents; (C-F) Pathway enrichment of the differential content of metabolites. The bubble size represented the number of enriched differential metabolites, and the gradient colour corresponded to the P-value of enrichment.

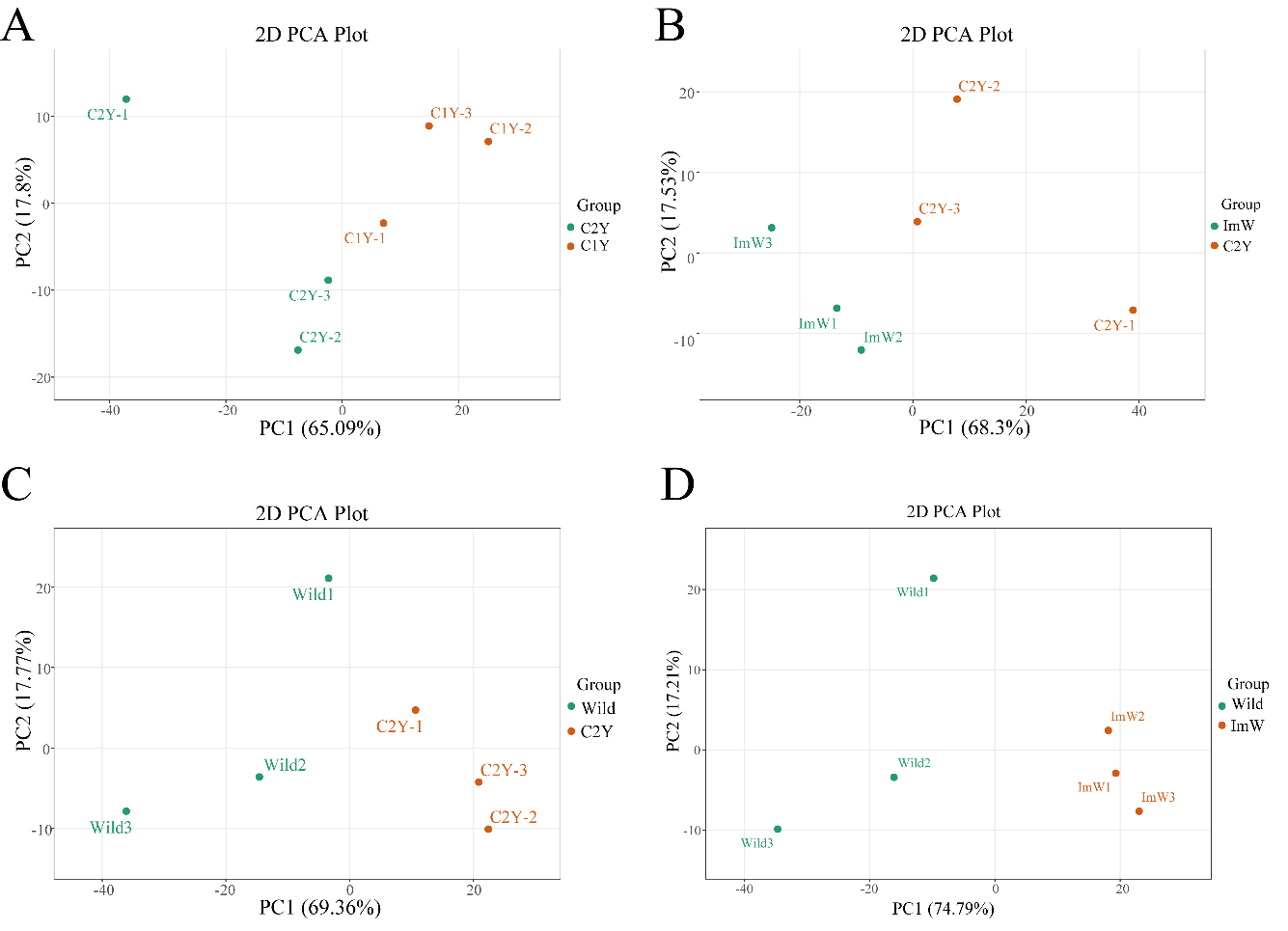

Fig. S2. PCA score plots showing metabolic clustering of pairwise comparison groups. (A) C1Y vs C2Y. (B) ImW vs C2Y. (C) Wild vs C2Y. (D) Wild vs ImW. Individual points represented biological replicates.

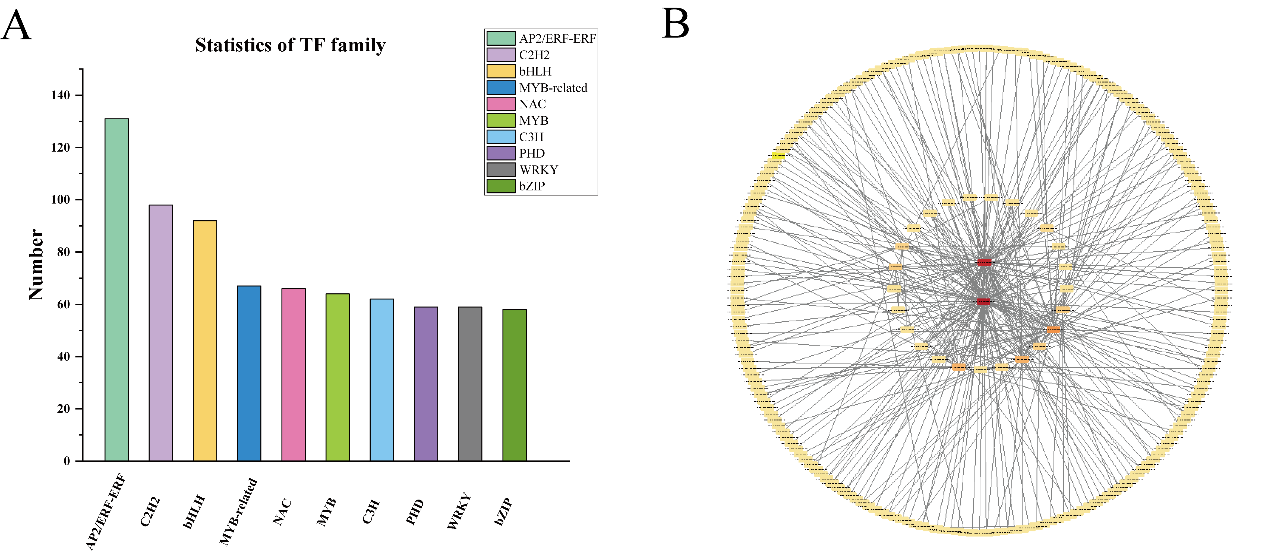

Fig. S3. Statistical analysis of transcription factor (TF) families and regulatory network of key genes in AMR. (A) The number of differentially expressed transcription factors belonging to the top 10 TF families; (B) Correlation regulatory network displayed the correlations between sesquiterpenoid biosynthesis-related differentially expressed genes and their upstream transcription factors.

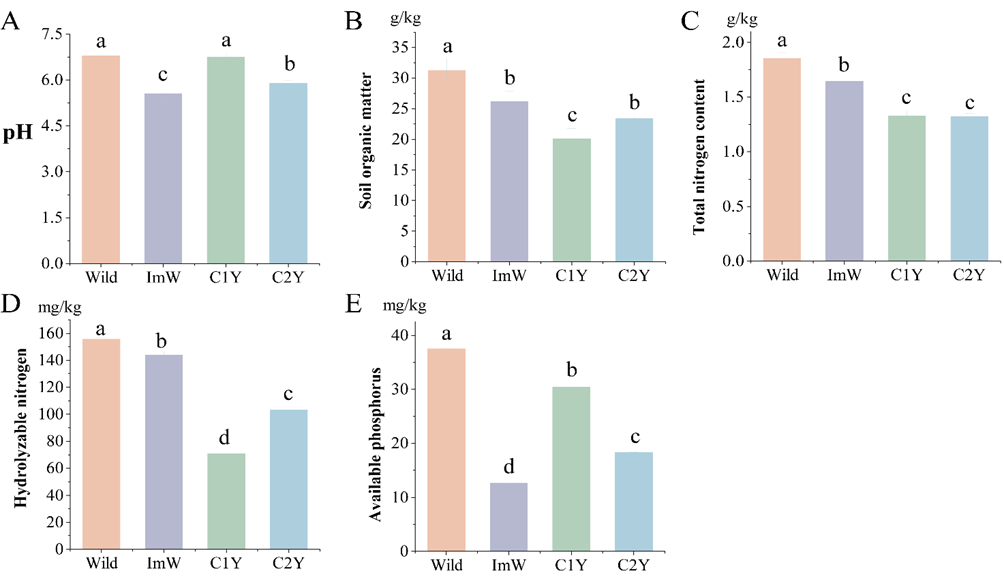

Fig. S4. Soil physicochemical properties across four AMR groups. (A) pH; (B) Soil organic matter; (C) Total nitrogen; (D) Hydrolyzable nitrogen; (E) Available phosphorus. Wild, ImW, C1Y and C2Y represented wild AMR, imitated wild AMR, one-year-old artificially cultivated AMR and two-year-old artificially cultivated AMR groups, respectively. Values were presented as mean ± SD (n = 3). Different lowercase letters indicated significant differences (*P* < 0.05).

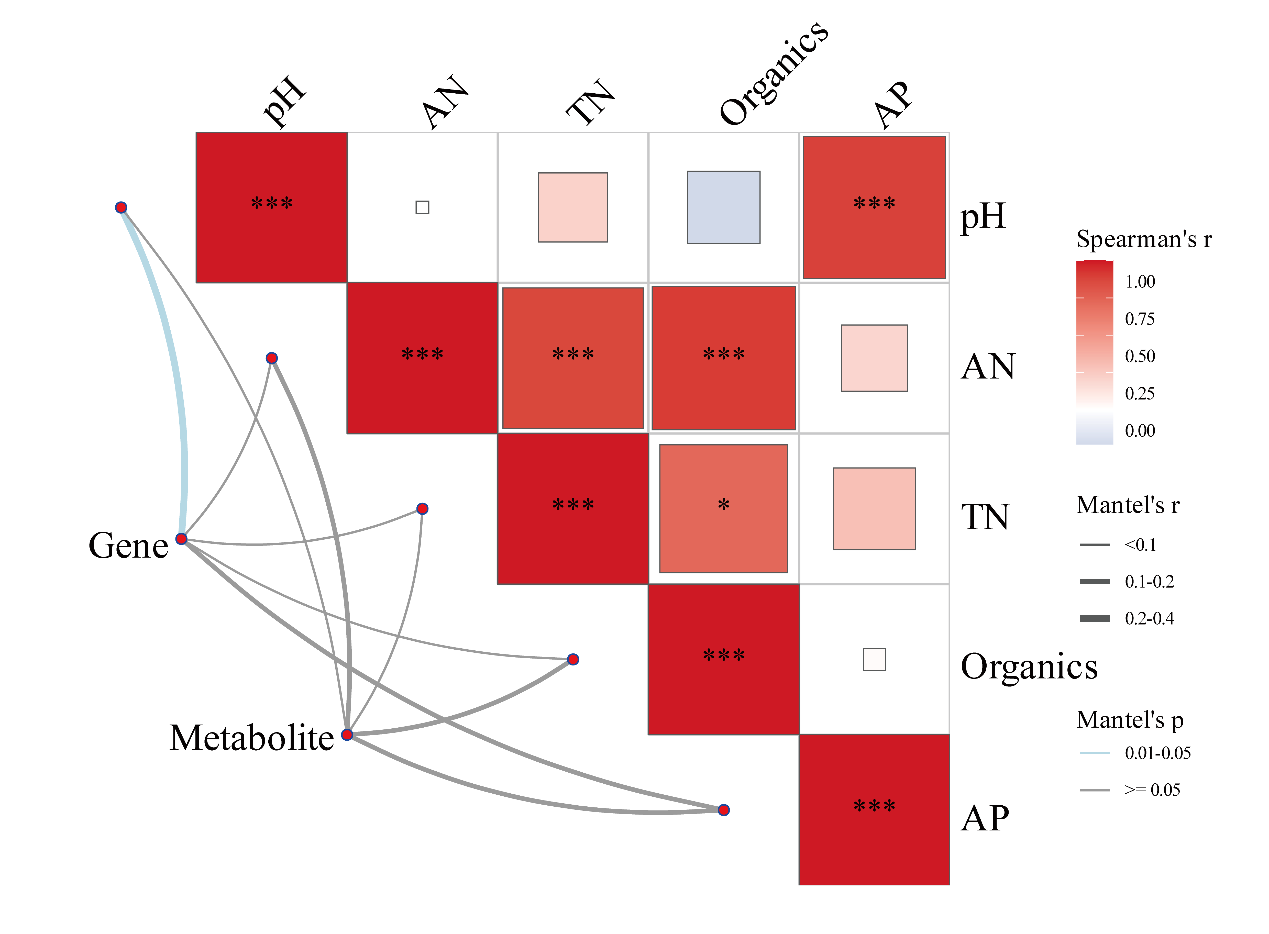

Fig. S5 Correlation heatmap and Mantel test visualization of differentially expressed genes, differential metabolites and soil edaphic factors. Heatmap colors represent Pearson correlation coefficients, while line thickness and color denote Mantel’s r and Mantel’s p values, respectively.

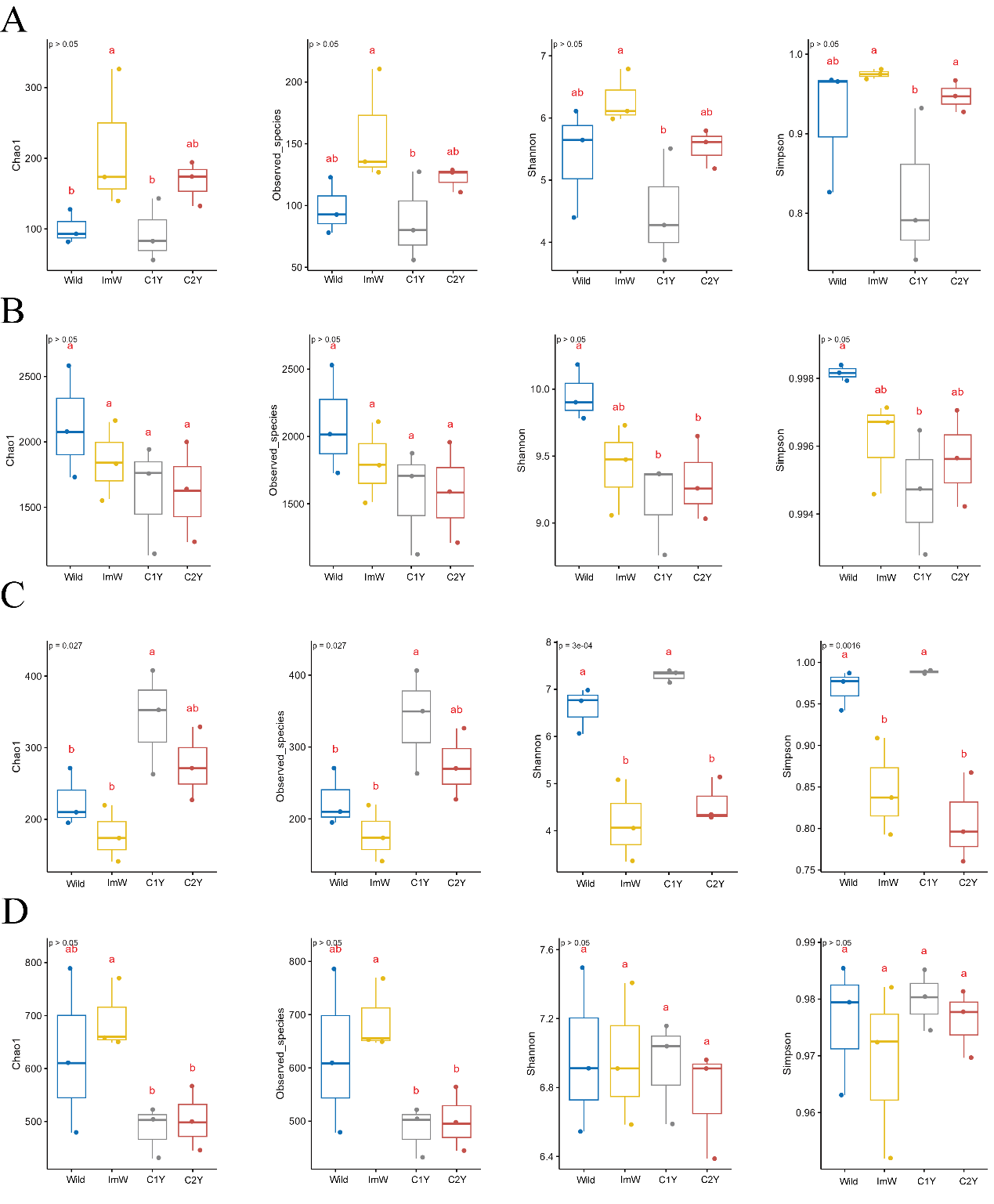

Fig. S6 Alpha diversity analysis of endophytic and rhizosphere microbial communities in AMR under different cultivation modes. (A) Alpha diversity indices (Chao1, Observed species, Shannon, Simpson) of endophytic bacterial communities; (B) Alpha diversity indices of rhizosphere bacterial communities; (C) Alpha diversity indices of endophytic fungal communities; (D) Alpha diversity indices of rhizosphere fungal communities. Groups labeled with different lowercase letters were significantly different (*P* < 0.05). Wild, ImW, C1Y and C2Y represented wild AMR, imitative wild-cultivated AMR, one-year artificially cultivated AMR, and two-year artificially cultivated AMR, respectively. The number of biological replicates was three.

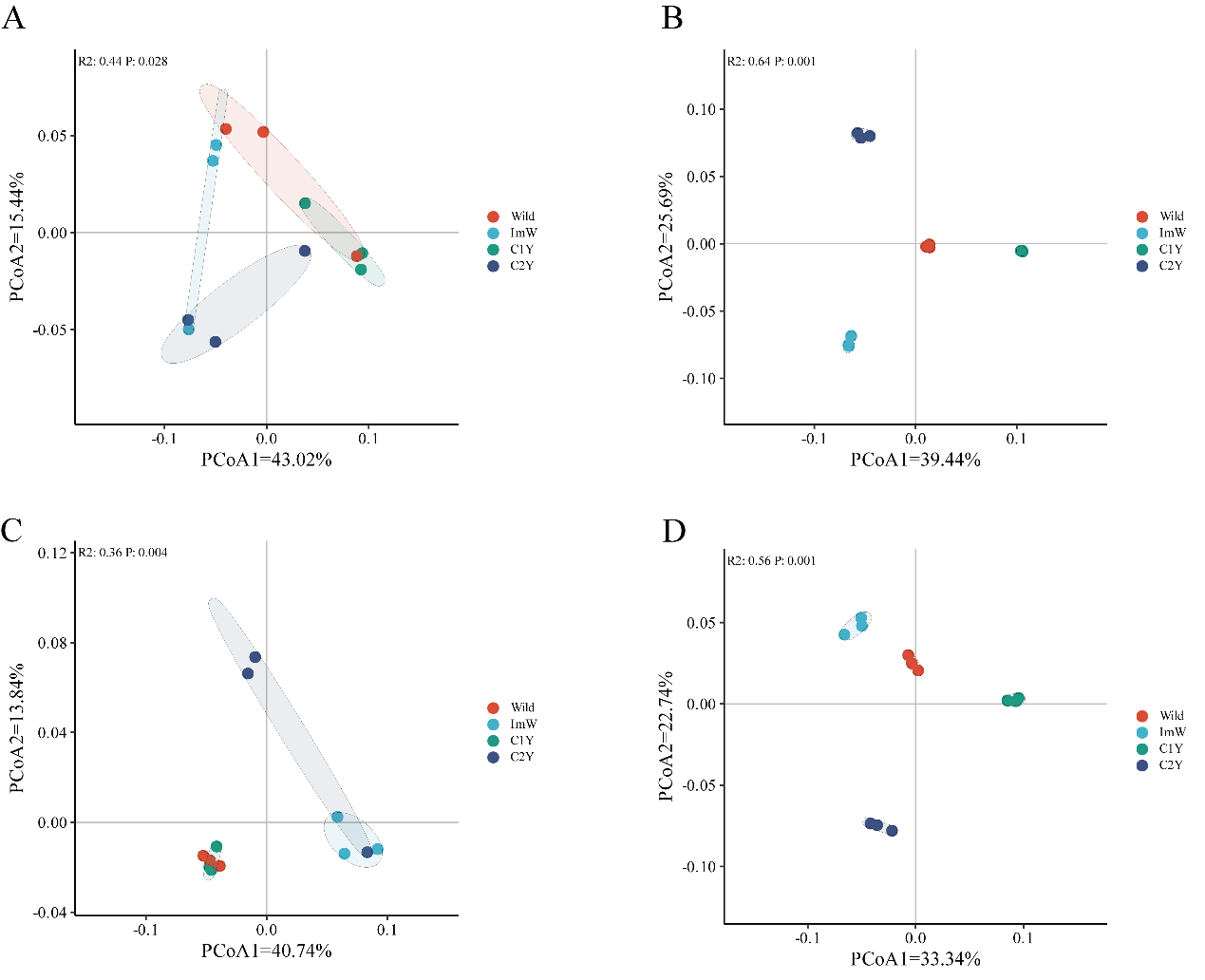

Fig. S7. Beta-diversity patterns of root microbial communities in AMR under different cultivation types. (A) PCoA of endophytic bacterial communities; (B) PCoA of rhizosphere bacterial communities; (C) PCoA of endophytic fungal communities; (C) PCoA of rhizosphere fungal communities. Wild, ImW, C1Y and C2Y represented wild AMR, imitative wild-cultivated AMR, one-year artificially cultivated AMR, and two-year artificially cultivated AMR, respectively. The number of biological replicates was three.

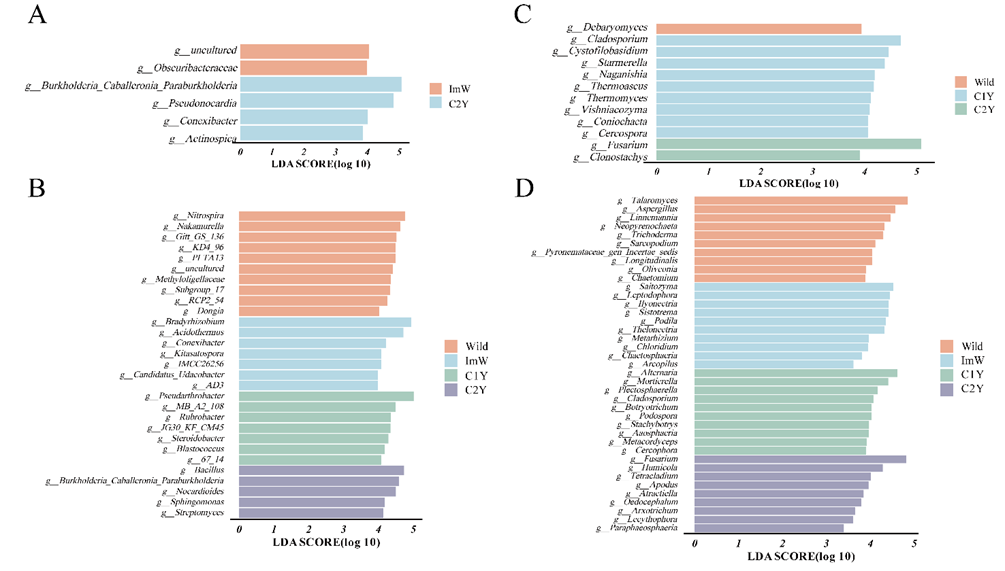

Fig. S8. Analysis of marker root-associated bacteria and fungi in AMR under different cultivation types. (A) LEfSe analysis of endophytic bacteria; (B) LEfSe analysis of rhizosphere bacteria; (C) LEfSe analysis of endophytic fungi; (D) LEfSe analysis of rhizosphere fungi. Wild, ImW, C1Y and C2Y represented wild AMR, imitative wild-cultivated AMR, one-year-old artificially cultivated AMR, and two-year-old artificially cultivated AMR, respectively. The number of biological replicates was three.
